# Identification of phospholipase Ds and phospholipid species involved in circadian clock alterations using CRISPR/Cas9-based multiplex editing of Arabidopsis

**DOI:** 10.1101/2024.01.09.574824

**Authors:** Sang-Chul Kim, Dmitri A. Nusinow, Xuemin Wang

**Affiliations:** Department of Biology, University of Missouri-St. Louis, St. Louis, MO 63121, USA; Donald Danforth Plant Science Center, St. Louis, MO 63132, USA

## Abstract

Reciprocal regulation between the circadian clock and lipid metabolism is emerging, but its mechanisms remain elusive. We reported that a lipid metabolite phosphatidic acid (PA) bound to the core clock transcription factors LATE ELONGATED HYPOCOTYL (LHY) and CIRCADIAN CLOCK ASSOCIATED1 (CCA1) and chemical suppression of phospholipase D (PLD)-catalyzed PA formation perturbed the clock in Arabidopsis. Here, we identified, among 12 members, specific PLDs critical to regulating clock function. We approached this using a multiplex CRISPR/Cas9 system to generate a library of plants bearing randomly mutated *PLDs,* then screening the mutants for altered rhythmic expression of *CCA1*. All *PLD*s, except for *β2*, were effectively edited, and the mutations were heritable. Screening of T2 plants identified some with an altered rhythm of *CCA1* expression, and this trait was observed in many of their progenies. Genotyping revealed that at least two of six *PLD*s (*α1, α3*, *γ1*, *δ*, *ε* and *ζ2*) were mutated in the clock-altered plants. Those plants also had reduced levels of PA molecular species that bound LHY and CCA1. This study identifies combinations of two or more PLDs and changes in particular phospholipid species involved in clock outputs and also suggests a functional redundancy of the six PLDs for regulating the plant circadian clock.

**One sentence summary:** This study identifies combinations of two or more phospholipase Ds involved in altering clock outputs and the specific phosphatidic acid species impacting the clock rhythms.

## Introduction

The circadian rhythms are 24-hour periodic changes in cellular and physiological processes driven by an internal timekeeping mechanism known as the circadian clock, enabling organisms to anticipate and adapt to the cyclic environment. A growing body of recent evidence has suggested that the biological clock is interconnected with and reciprocally regulated by metabolism. While the clock regulates temporal metabolic activities by modulating the daily abundance of cellular factors involved in metabolism, some metabolites serve to provide input to the clock (Delezie and Challet, 2011; Pan et al., 2020; Grosjean et al., 2023). Despite the emerging pathways through which metabolic signals influence the plant circadian system (Buckley et al., 2023), however, mechanisms of how lipid metabolism affects the clock function are poorly understood in plants, mainly due to the scarce knowledge about molecular interactions between lipid metabolites and clock components. Recently, we reported that a vital lipid mediator phosphatidic acid (PA) bound to and suppressed the ability of the core clock transcription factor LATE ELONGATED HYPOCOTYL (LHY) and its closely related CIRCADIAN CLOCK ASSOCIATED1 (CCA1) to bind the promoter of their target genes (Kim et al., 2019). We also found that cellular PA abundance directly correlated with the period of circadian rhythms, such as the vertical movement of plant leaves and the oscillatory expression of the LHY/CCA1 target *TIMING OF CAB EXPRESSION1* (*TOC1*) (Kim et al., 2019). In addition, our recent results indicate that the PA-clock interaction attenuates the LHY/CCA1 promotion of fatty synthesis and storage lipid accumulation (Kim et al., 2023). These findings suggest that PA is a vital regulator of the plant circadian clock. Furthermore, given that PA is the pivotal metabolic intermediate formed during the conversion among many other lipids, the PA-LHY/CCA1 interaction may function as a cellular conduit to integrate lipid metabolism with the clock in plants.

The cellular level of PA is highly dynamic and tightly regulated by diverse enzymes that catalyze its formation and degradation (Kim and Wang, 2020). Multiple gene families encode the PA-producing and removing enzymes. Many display different lipid substrate preferences, catalytic requirements, and/or subcellular distribution, leading to the high complexity of PA metabolic routes (Yao et al., 2024). Phospholipase D (PLD) hydrolyzes common membrane phospholipids, such as phosphatidylcholine (PC) and phosphatidylethanolamine (PE), to produce PA by releasing their alcohol moiety. Arabidopsis PLD family comprises 12 members, designated as α1/2/3, β1/2, γ1/2/3, δ, ε, and ζ1/2, classified according to amino acid sequence and enzymatic properties (Qin and Wang, 2002). They can be activated differently in response to diverse stimuli, and these differences account for the highly dynamic nature of PA contents and compositions depending on environmental conditions (Hong et al., 2016; Ali et al., 2022; Deepika and Singh, 2022; Kolesnikov et al., 2022). This is evidenced by, in addition to early studies, recent reports showing a rapid accumulation of PA by PLDα1 in response to magnesium stress and rhizobium infection, β2 during hypoxia, δ under heat and osmotic stress, and ε and ζ2 in nitrogen and phosphate deficit, respectively, with a few specific PA molecular species contributing to the accumulation of total PA in some cases (Su et al., 2018; Premkumar et al., 2019; Song et al., 2020; Kocourková et al., 2021; Liu et al., 2021; Zhang et al., 2021; Kim et al., 2022; Yao et al., 2022 a and b). Not only for plants to cope with unfavorable conditions, PLDs are also required for normal physiological processes mediated by their direct interaction with effector proteins, regulation of cytoskeletal dynamics and vesicle trafficking, and membrane remodeling and degradation (Hong et al., 2016; Deepika and Singh, 2022).

All PLDs can use primary alcohols as a substrate to produce phosphatidylalcohol instead of PA, and 1-butanol has often been used to suppress PLD-catalyzed PA formation in plants (Munnik et al., 1995). We previously found ∼40% reduction of basal PA levels in Arabidopsis treated with 1-butanol, which shortened the circadian rhythm period (Kim et al., 2019). Since PLDs play different roles in stress responses, it is conceivable that specific PLD(s) produces the cellular pool of PA that interacts with LHY/CCA, and the reduction of the pool size leads to dissonance between the internal and external rhythms and eventually compromises plant response to stress. Thus, identifying the PLD(s) involved in this process is critical to determine how lipid metabolism communicates with the clock components to affect the circadian rhythms and stress responses. However, our preliminary data indicated that single T-DNA mutants of individual *PLD*s had no significant effect on the clock, indicative of the functional redundancy of PLDs for the clock regulation. Moreover, LHY and CCA1 only bound to specific PA species, and neither their levels in Arabidopsis nor LHY-binding to its targets significantly changed in a double knockout mutant of the most abundant PLDs, *α1* and *δ* (Kim et al., 2019). This implies that other PLDs besides the two major ones are required to form LHY/CCA1-binding PA species. Thus, here, we generated and screened Arabidopsis plants with various combinations of *PLD*s randomly mutated by a multiplex CRISPR/Cas9 system. In this approach, we successfully produced a library of plants with mutations of different *PLD*s and identified several *PLDs* essential for circadian clock function. In addition, lipidomic analysis of those mutants identified specific PA species associated with clock alterations.

## Results

### Generation of a multiplex CRISPR/Cas9-edited PLD mutant library

To target the 12 *PLD*s by multiplex CRISPR system, we used a toolkit that allows the assembly of nuclease constructs expressing up to 32 single guide RNAs (sgRNAs) in a single nuclease vector (Ordon et al., 2017; Grutzner et al., 2021; Stuttmann et al., 2021). This toolkit contains Dicot Genome Editing (pDGE) vector system that is composed of ‘shuttle’ vectors for the preparation of sgRNA transcriptional units, intermediate ‘multi-multi’ vectors for assembly of more than 8 sgRNAs, and pre-assembled ‘recipient’ vectors with *Cas9* (Figure 1A). We targeted two different sites in each *PLD* for enhanced editing efficiency (a total of 24 sgRNAs; Figure S1). Eight hybridized oligonucleotides for sgRNAs were first cloned into 8 different shuttle vectors (pDGE332-342) for each of three sets of the 8 oligonucleotides (1-8, 9-16 and 17-24 in Figure 1A). Since all the shuttle vectors contain two *BamHI* sites, and one of them is within the cloning cassette to be replaced by the oligonucleotides, successful cloning can be verified by cutting the DNA constructs with *BamHI*: a large DNA fragment resulting from the cloned constructs (∼3 kb) and an additional smaller fragment from empty vectors (∼0.8-0.9 kb) (Figure 1B). The three sets of the shuttle vector constructs were then subcloned into three different multi-multi vectors (pDGE503-505 in Figure 1A). *BpiI* digestion of the multi-multi vector constructs from 5 randomly selected *E. coli* clones resulted in the correct size of the insert DNA (∼1.7 kb; the 8 sgRNA transcriptional units with some flanking nucleotides) from at least one clone for each of the three constructs (Figure 1C). The three confirmed multi-multi vector constructs were finally cloned into the recipient vector (pDGE668 in Figure 1A). Three of four randomly selected *E. coli* clones had the correct size of the insert DNA (5,007 bps; the 24 sgRNA transcriptional units with some flanking nucleotides), as revealed by *HindIII* digestion (Figure 1D). The final confirmed recipient vector construct was transformed into Arabidopsis for editing its *PLD*s.

**Figure 1.**
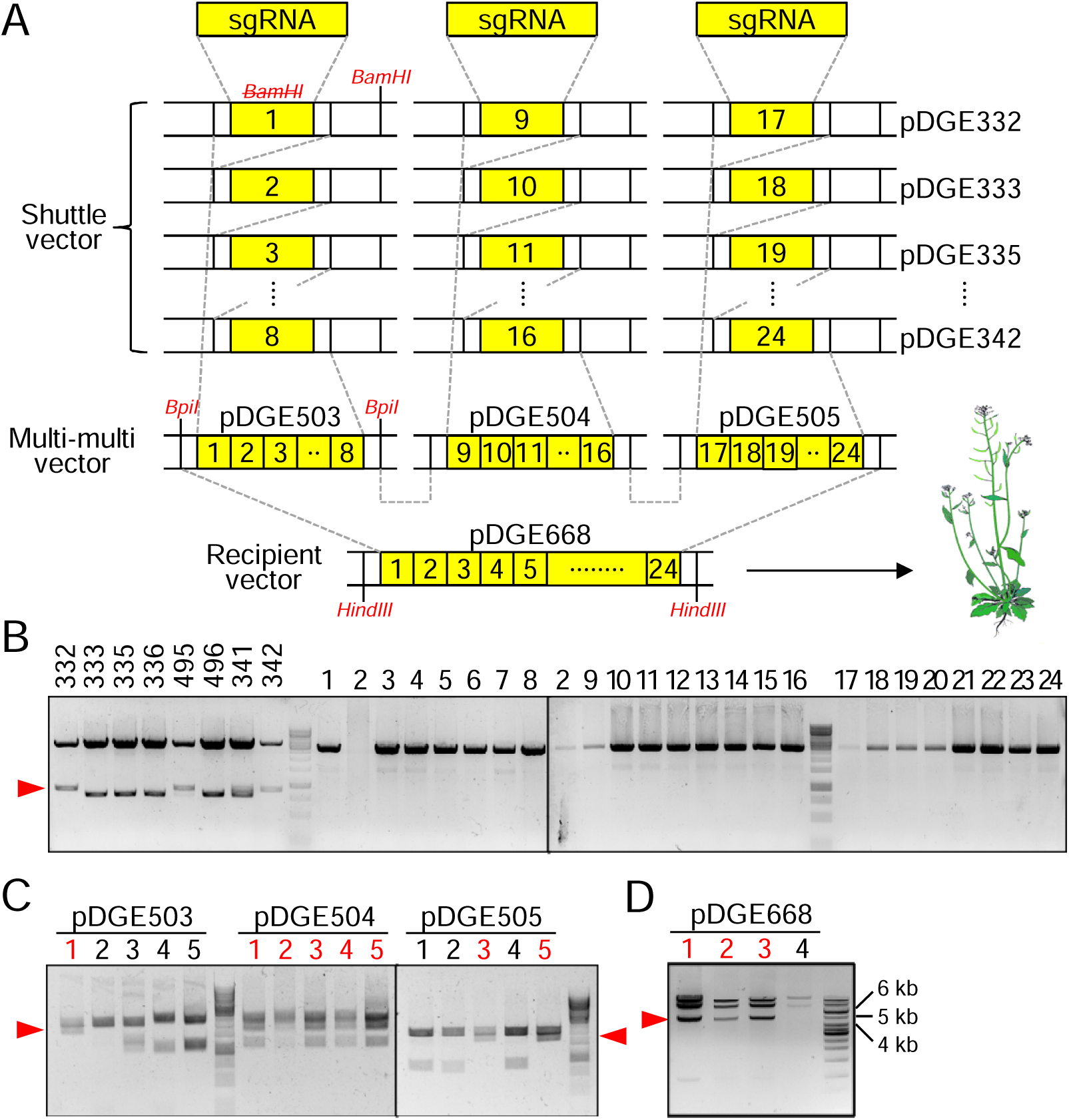
Cloning of oligonucleotides for 24 sgRNAs targeting 12 *PLD*s. **A.** Schematic overview of the cloning procedure. 24 sgRNAs (two target sites for each *PLD*) were cloned sequentially in shuttle vectors, multi-multi vectors, and finally recipient vector for Arabidopsis transformation, in order by Golden Gate cloning. Two cohesive ligation sites are connected with dashed lines, and restriction enzyme sites to confirm the cloning are in red (*BamHI* site removed by sgRNA is struck through). **B.** Confirmation of shuttle vector constructs. Empty vectors (332-342) and cloned DNA constructs (1-24) were cut with *BamHI*. Arrowhead indicates the position of the small fragment resulted from *BamHI* cut in the empty vectors. Note that ‘2’ was failed first (left lane) and retried later (right lane). **C.** Confirmation of multi-multi vector constructs. Plasmids from 5 random *E. coli* clones were cut with *BpiI* for each multi-multi vector. Arrowhead indicates the position of the fragment containing all 8 sgRNAs. **D.** Confirmation of recipient vector constructs. Plasmids from 4 random *E. coli* clones were cut with *HindIII.* Arrowhead indicates the position of the fragment containing all 24 sgRNAs. For C and D, the clone numbers with the desired fragments are highlighted in red.

The recipient vector contains a selection marker against kanamycin, and successful plant transformation was verified by PCR amplification of *Cas9* in the genomic DNA of 10 kanamycin-resistant T1 plants (Figure 2A). To see whether and which *PLD*s were edited in the transformants, we performed PCR to amplify a genomic DNA region spanning the two target sites for each *PLD* in six independent *Cas9*-positive plants. We checked the PCR products on a gel to detect large insertion, deletion or chimeric mutations that the natural non-homologous end-joining might have introduced. For all *PLD*s except for *β2* and *ε*, some of the plants had one or more PCR products smaller than and in addition to the one observed in wild-type (WT), indicative of heterozygous deletion mutations introduced (Figure 2B; the exact size of PCR products from WT in Figure 2C). PCR results from six additional transformants (a total of 12 individual plants) are shown in a diagram in Figure 2D. At least three plants had a mutation in each *PLD* (>25% frequency), with a notably high frequency of *γ3* (100%), and no mutation was found in *β2* and *ε* (Figure 2D). In each plant, three up to nine *PLD*s were mutated, and no plant appeared to have all *PLD*s mutated (Figure 2D). It should be noted that these data are based on visual inspection of the DNA bands, and thus, the actual mutation frequency is expected to be higher than that revealed by the PCR/gel-based analyses. The results here indicate that our multiplex CRISPR/Cas9 system effectively edited at least 10 PLDs.

**Figure 2.**
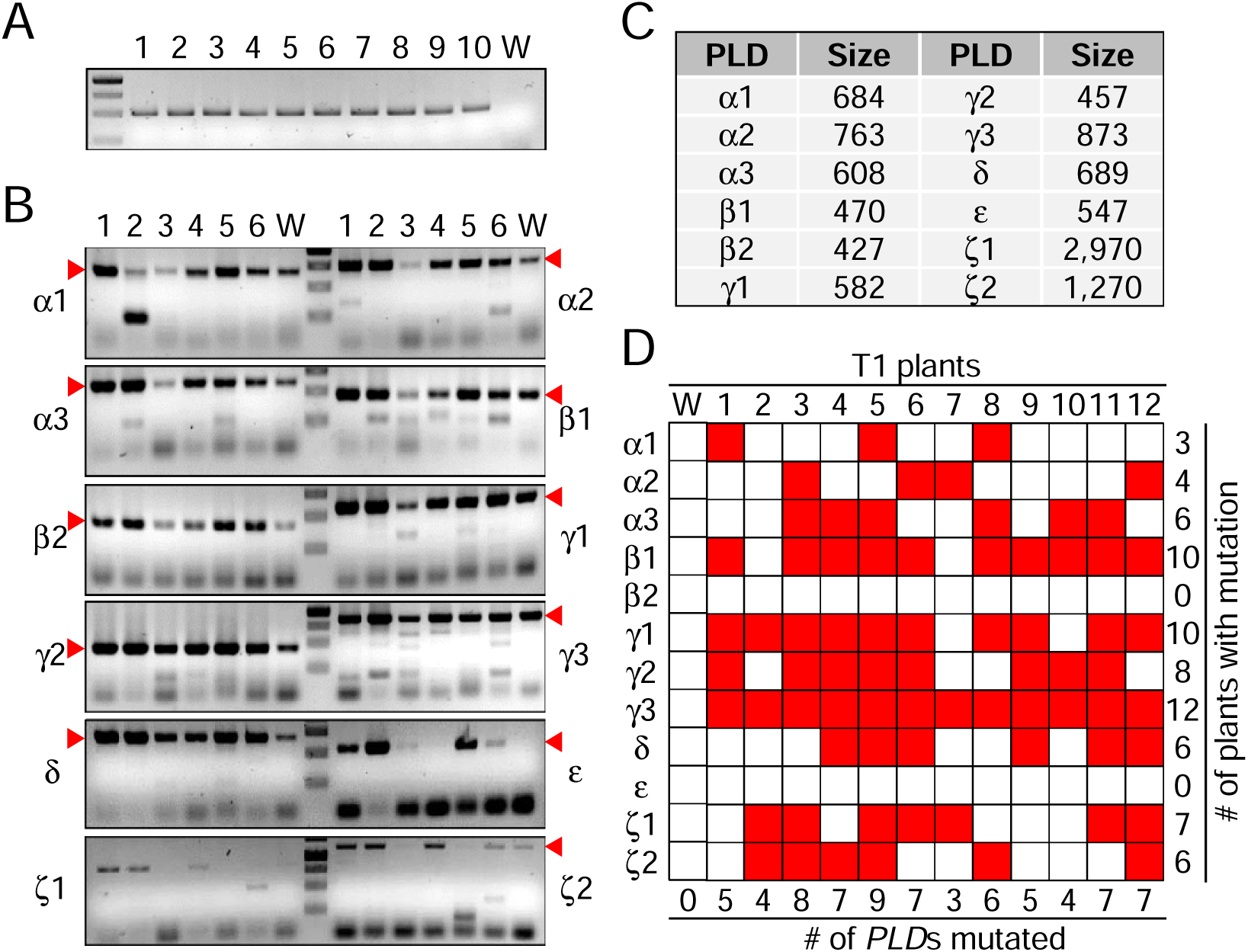
Detection of deletion mutations in T1 plants. **A.** Presence of *Cas9* in kanamycin-resistant plants. Genomic DNA was extracted from 10 kanamycin-resistant T1 plants, and PCR was performed using *Cas9* primers. W, wild-type. **B.** PCR detection of the deletion mutation. PCR was performed using *PLD* primers from genomic DNA in 6 *Cas9*-positive T1 plants. Arrow head indicates the position of PCR product from wild-type (unmutated) allele. Note that wild-type *PLD*ζ*1* could not be amplified due to its large size and is not shown here. Size of marker DNA (middle lane) is 250, 500, 750, and 1000 bps from the bottom. W, wild-type. **C.** Size (bps) of PCR product from wild-type (unmutated) *PLD*. **D.** Frequency of deletion mutations in each *PLD*. Deletion mutations in 12 individual T1 plants analyzed as in B are marked in red. The numbers of *PLD*s mutated in each plant and plants with mutation in each *PLD* are at the bottom and on the right, respectively.

### Detection of mutations in T2 plants and sequencing analysis of selected PLDs

The vector system provides a gene encoding red fluorescence protein (RFP) expressed under the seed-specific promoter *OLE1p* as a counter-selection marker for non-transgenic seeds. We harvested and combined seeds from the 12 T1 transformants and selected the seeds without red fluorescence to obtain non-transgenic (*Cas9*-free) T2 plants to avoid novel somatic mutations caused by additional edits. Approximately a quarter of the screened T2 seeds exhibited no red fluorescence following the classic Mendelian model of single insertion (Figure 3A). The absence of the transgene in the selected seeds was further confirmed by the failure of PCR amplification of *Cas9* in the genomic DNA of their seedlings (Figure 3B). We then performed the same PCR analysis to evaluate T1 plants for each *PLD* in 10 *Cas9*-negative T2 plants. As expected, the overall mutation frequency is much lower, and some plants had a mixture of homozygous and heterozygous mutations, as demonstrated by a single smaller PCR fragment without the one from WT (Figure 3C). Notably, some *PLD*s produced two different small fragments and no WT fragment, indicative of the ‘biallelic’ heterozygous mutation with the two alleles mutated differently (Figure 3C). Among the 10 plants, nine had a homozygous or heterozygous deletion mutation in one to three *PLD*s, and each *PLD* was mutated in one to three plants, again except for *β2* and *ε* that were unmutated in all plants tested (Figure 3D).

**Figure 3.**
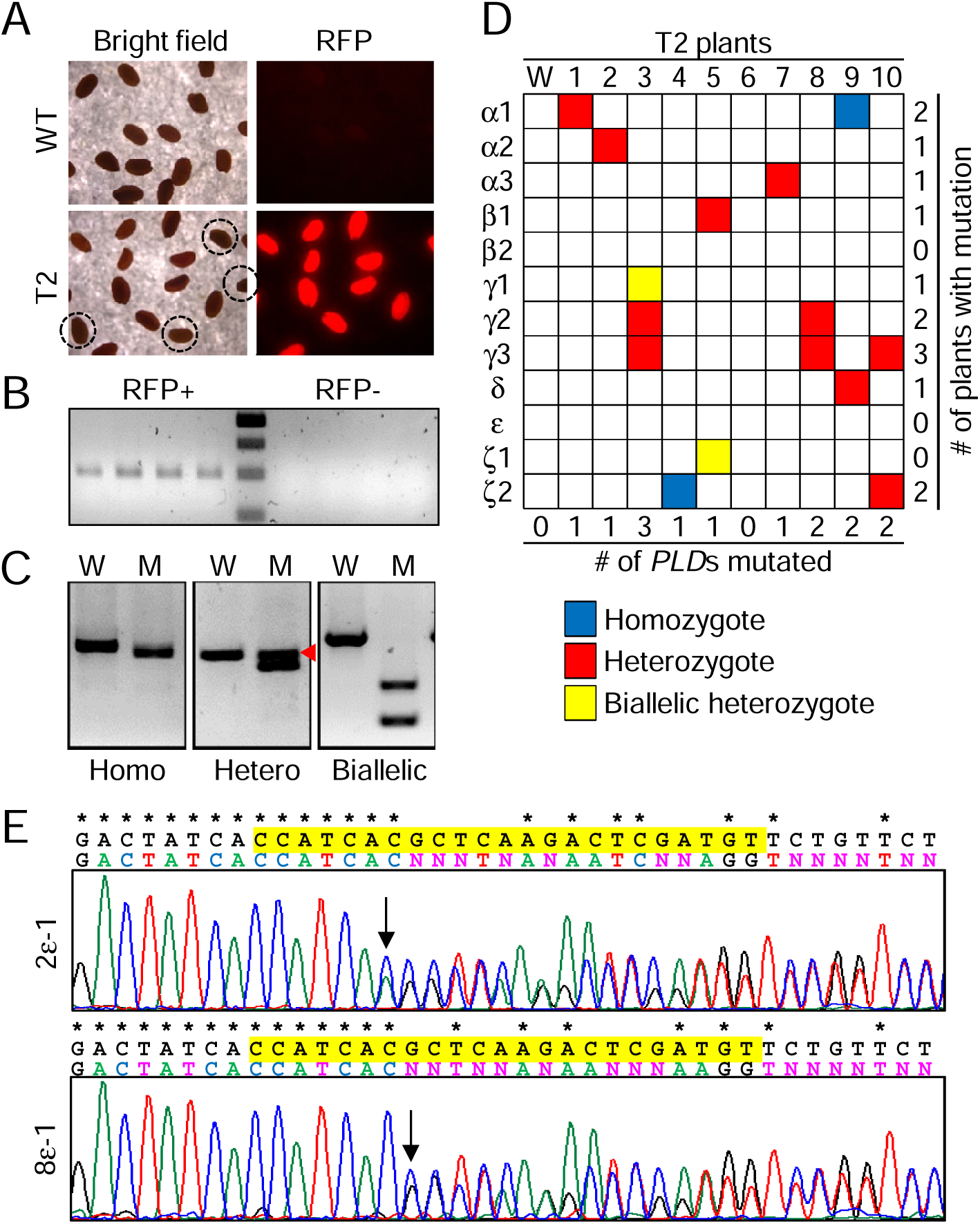
Detection of deletion mutations in T2 plants. **A.** Selection of non-transgenic T2 seeds. T2 seeds were observed under a fluorescence microscope with RFP filter. RFP-negative seeds are circled in the bright field image. **B.** Absence of *Cas9* in RFP-negative plants. Genomic DNA was extracted from randomly selected RFP-positive and negative T2 plants. PCR was performed using *Cas9* primers. **C.** PCR detection of various zygosities in deletion mutation. PCR was performed using *PLD* primers from genomic DNA in non-transgenic T2 plants. Shown here are representative images showing homozygote, heterozygote, and biallelic heterozygote. Arrowhead indicates the PCR product from wild-type (unmutated) allele. W, wild-type; M, mutant (*PLD*-altered). **D.** Frequency of deletion mutations in each *PLD*. Deletion mutations in 10 individual non-transgenic T2 plants analyzed as in C are color-coded for their zygosities. The numbers of *PLD*s mutated in each plant and plants with mutation in each *PLD* are at the bottom and on the right, respectively. **E.** Sequencing chromatograms for *PLD*ε. Genomic DNA was extracted, and *PLD*ε was PCR-amplified and sequenced. Plant number, PLD gene and target site (1 or 2) are indicated on the left. Above the chromatograms, wild-type (top) and mutant (sequenced; bottom) sequences are aligned with asterisks for the identical nucleotides, and Cas9 target sequence in WT is in yellow. Arrow indicates the first overlapping peaks.

Since *PLDβ2* and *ε* appeared to have no mutation in both T1 and T2 plants, we carried out a more comprehensive analysis of 6 of the 10 T2 plants by amplicon sequencing of the two *PLD*s to identify their mutations unable to be detected by the gel-based method. We still found no mutation in *PLDβ*2 at both target sites in the 6 plants, indicating its very low efficiency for editing. However, two plants had a heterozygous insertion or deletion mutation in *PLDε* at the first target site, as identified in the sequencing chromatograms by a series of single peaks followed by two overlapping peaks continuing to the end (Figure 3E). The data here indicate that most of our non-transgenic T2 plants have a mutation, either homozygous or heterozygous, in at least one of the 12 *PLD*s, except for *β2*.

### Screening of the genome-edited PLD library to select clock-altered mutant plants

Based on our previous finding that inhibition of PLD-catalyzed PA formation shortened the period of *TOC1* expression (Kim et al., 2019), we screened the *Cas9*-negative T2 plants pre-transformed with luciferase reporter gene expressed by *TOC1* or *CCA1* promoter (*TOC1::LUC* or *CCA1::LUC*) to select the ones with a shorter period of the clock gene expression. PCR analyses confirmed the absence of *Cas9* and the presence of *LUC* in six RFP-negative T2 plants randomly chosen for each of the two *LUC* lines (Figure 4A). The luciferase activity was monitored over time to measure the rhythmic patterns of *CCA1* and *TOC1* expression in T2 seedlings. Among several plants displaying clear rhythmicity of the reporter activity, four with *CCA1::LUC* and two with *TOC1::LUC* showed a shorter period when compared to their respective WTs (Figure 4B). We then measured the period of *CCA1* expression in ∼20 progenies (T3) for each of the four T2 plants showing altered *CCA1* expression. Interestingly, most T3 plants appeared to have a shorter period of *CCA1* expression than WT, largely regardless of their parental lines. Figure 4C shows two representative plants for each family displaying the shortest period. Computational period estimation (Zieliński et al., 2021) revealed that *CCA1* expression in WT was nearly 24-hour periodic (24.12 h), and the T3 plants had an average of 23.29-23.62 hours and differed from WT by up to 2.46 hours (Figure 4D). Our data suggest that the phenotypic change in the clock gene expression is heritable and likely caused by the CRISPR-induced *PLD* mutations.

**Figure 4.**
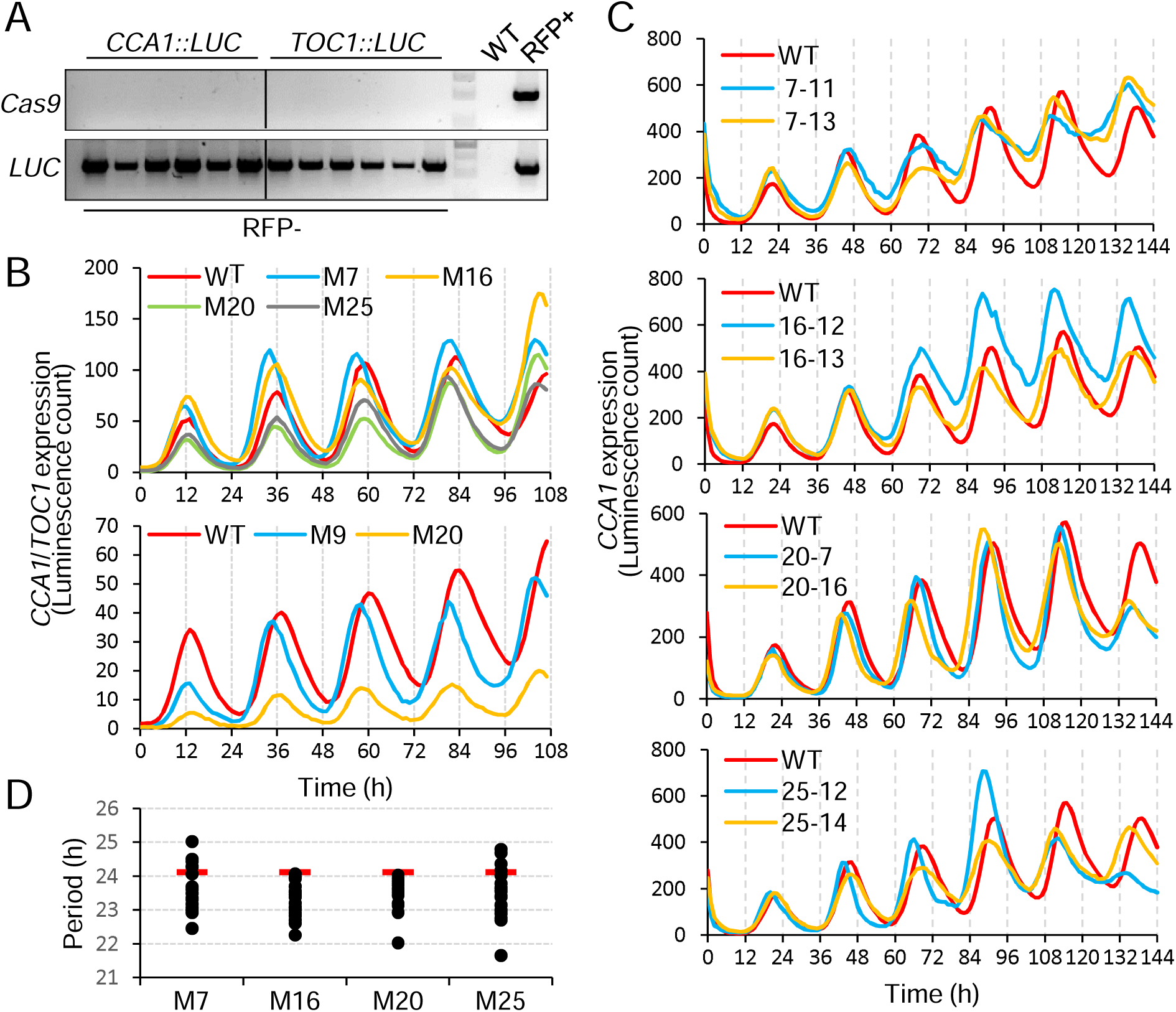
Rhythmic patterns of *CCA1* and *TOC1* expression. **A.** Absence of *Cas9* and presence of *luciferase* in RFP-negative plants. Genomic DNA was extracted from randomly selected RFP-negative T2 plants with *CCA1/TOC1::LUC*. PCR was performed using *Cas9* and *luciferase* (*LUC*) primers. Wild-type (WT) and RFP-positive (RFP+) plants were included as controls. **B.** *CCA1*/*TOC1* expression patterns in *Cas9*-negative T2 plants. Luciferase activity driven by *CCA1* (top) or *TOC1* (bottom) promoter was monitored over time. Shown here are the plants with a shorter period of *CCA1*/*TOC1* expression than WT. Note that plants with *CCA1* and *TOC1* were grown with 12-hour difference for convenience of data analysis, appearing in phase with each other. WT, wild-type; M, mutant (*PLD*-altered). **C.** *CCA* expression patterns in T3 plants. The 4 T2 plants in B were grown to the next generation and their T3 plants were measured for luciferase activity as in B. Two representative plants with the shortest period of *CCA1* expression are shown here for each family (M7, M16, M20, and M25 from the top). **D.** Periods of *CCA1* expression in T3 plants. Period was estimated using BioDare2 with linear detrend and FFT-NLLS method. Red bar indicates wild-type period.

### Identification of specific PLDs involved in altered CCA1 expression

We genotyped the 8 T3 plants shown in Figure 4C by amplicon sequencing to determine which PLDs were mutated in the clock-altered plants. All *PLD*s in WT were intact, as expected. Some of the selected plants had homozygous mutations in *PLDα1* and *γ1* at either or both target sites, and all were identified as having an insertion or deletion of one nucleotide (Figure 5A). All homozygous mutations in the first target site of *PLDα1* had a deletion of an ‘A’ at the same position (sixth nucleotide in the target site; Figure 5A). We also found heterozygous mutations in six *PLD*s (*α1*, *α3*, *γ1*, *δ*, *ε* and *ζ2*), as identified by the overlapping peaks extending in the chromatograms as in Figure 3E, and the other six (*α2*, *β1*, *β2*, *γ2*, *γ3* and *ζ1*) were unmutated in all plants (Figure 5B). Combinatorial analyses of the overlapping peaks revealed that all heterozygous mutations, like homozygotes, were a frame-shift mutation and not biallelic; *PLDα1* in plant 25-14 and *γ1* in 16-12 turned out to be mutated in both target sites in the same chromosome (Figure 5B). *PLDα1* in 25-12 and 25-14 had both a deletion and insertion of one nucleotide, but the deletion mutation resulted in a premature stop codon at the 37th amino acid (Figure 5C for 25-14). All 8 plants had a homozygous or heterozygous mutation in at least two *PLD*s, with the equally highest frequency of *PLDα1*, *α3,* and *γ1* (four plants) followed by *ε* and *ζ2* (two plants), and then *δ* mutated in only one plant (Figure 5B). The identification of the mutated *PLD*s here narrows down the number of candidates likely involved in the clock function and indicates that the six PLDs may play a redundant role in the proper functioning of the circadian clock.

**Figure 5.**
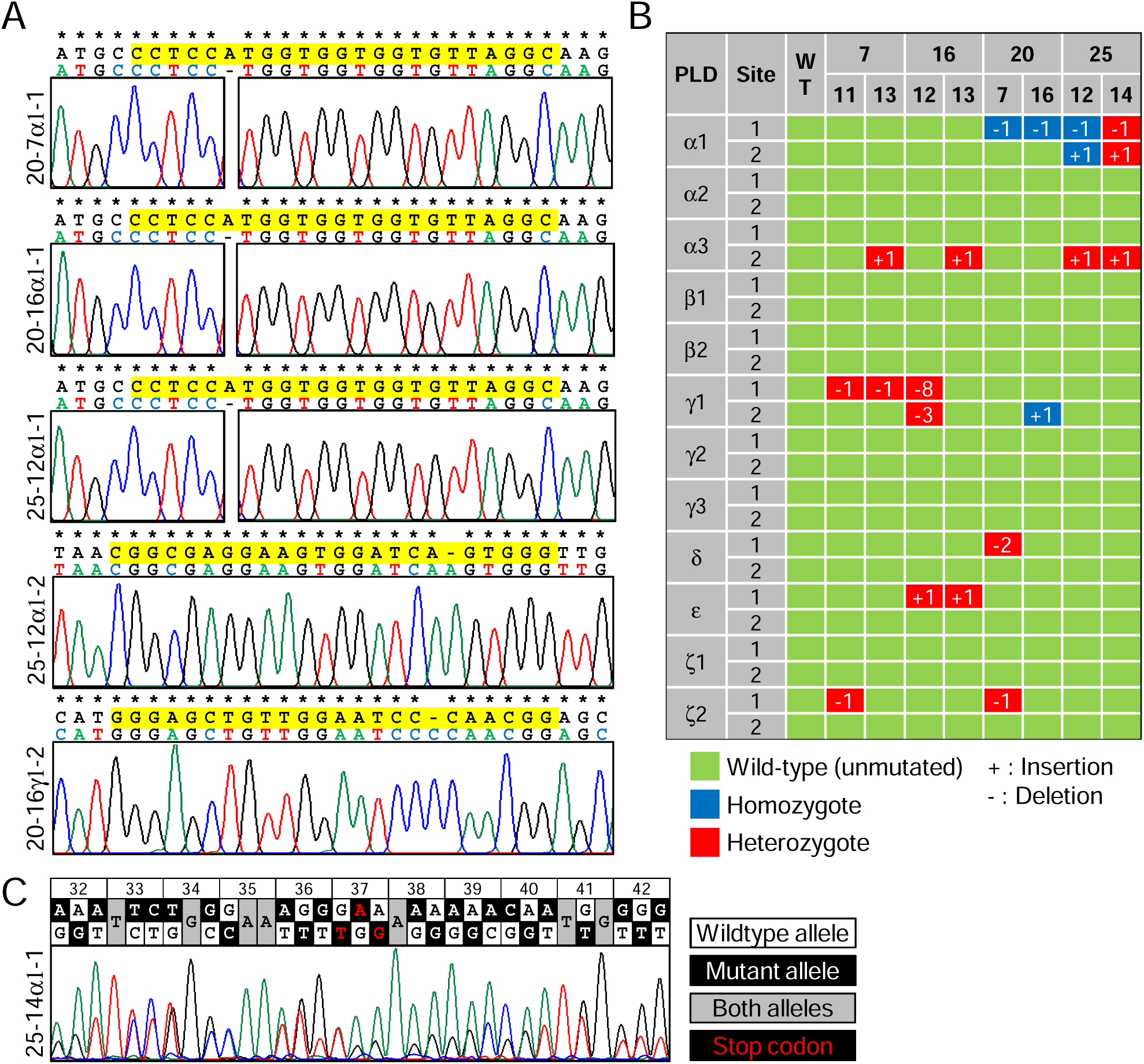
Genotyping of selected T3 plants by sequencing. **A.** Sequencing chromatograms for homozygous mutations. Genomic DNA was extracted, and *PLDs* were PCR-amplified and sequenced. Plant number, PLD gene and target site (1 or 2) are indicated on the left. Above the chromatograms, wild-type (top) and mutant (sequenced; bottom) sequences are aligned with asterisks for the identical nucleotides, and Cas9 target sequence in WT is in yellow. **B.** Genotype of the selected T3 plants. All *PLD*s in the 8 T3 plants are color-coded for their zygosities, with the number of nucleotides mutated. In the ‘site’ column, 1 and 2 indicate the first and second target sites, respectively. In the plant number rows, top and bottom rows are T2 and T3 plants, respectively. WT, wild-type. **C.** Sequencing chromatogram for 25-14α1-1. Two nucleotides at each peak position are shown in black and white, with the position number of amino acid for each codon at the top. The target site is upstream and not shown here.

### PLDα protein levels in the T3 plants with altered CCA1 expression

It was intriguing that the clock phenotype was altered by heterozygous and homozygous mutations of *PLD*s, suggesting that the phenotype was likely caused by a protein dosage effect of PLD. Thus, we carried out immunoblotting to analyze PLD protein abundance in all *PLDα*-mutated T3 plants (7-13, 16-13, 20-7, 20-16, 25-12, and 25-14) using our available PLDα-specific antibody (Hong et al., 2008b). Probing cytosolic glyceraldehyde-3-phosphate dehydrogenase (GAPC) verified the equal amounts of samples examined across the different plants (Figure 6A). PLDα was detected in WT and the plants with *PLDα1* unmutated (7-13 and 16-13), but not when *PLDα1* was homozygously mutated (20-7, 20-16, and 25-12 in Figure 6A). This was regardless of *PLDα2* and *α3* mutations, probably due to the much lower abundance of α2 and α3 proteins (Hong et al., 2016). Interestingly, PLDα was also undetectable in the plant with a heterozygous mutation in *PLDα1* (25-14 in Figure 6A). This explains why the plants with heterozygous mutations, at least in *PLDα1*, exhibited the altered clock phenotype. However, it is unclear how the heterozygous mutation leads to no detectable protein expression. Further interpretation of the genotyping and protein data will follow in the Discussion.

**Figure 6.**
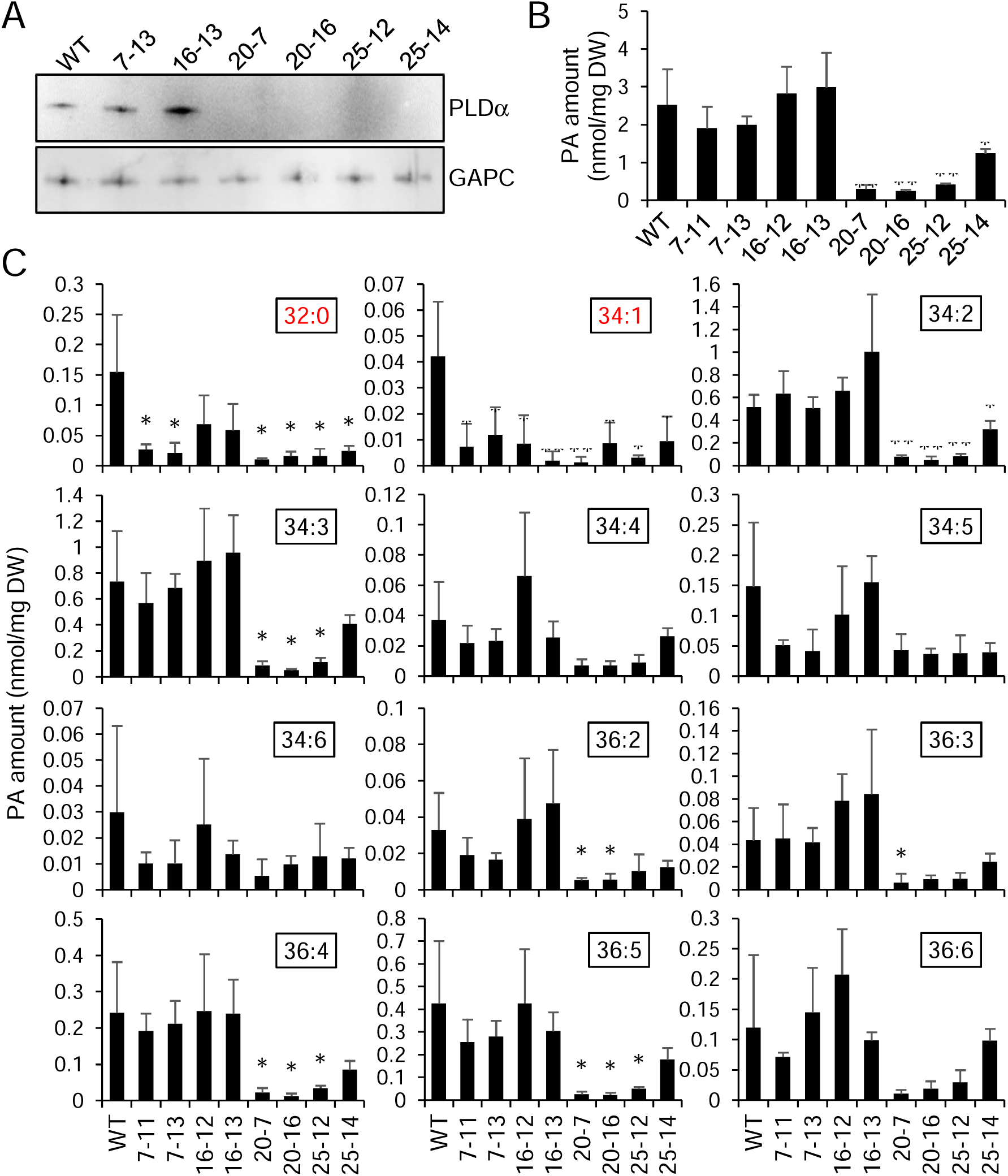
Protein abundance of PLDα and PA levels in the selected T3 mutants. **A.** Protein abundance of PLDα. Total proteins were extracted from the *PLDα*-mutated T3 plants, and PLDα was probed by immunoblotting with an anti-PLDα antibody raised against castor bean PLDα (top). Anti-GAPC antibody was used as a loading control (bottom). **B.** Total PA levels. Total lipids were extracted from the selected T3 mutants, and PA was separated and quantified by ESI-MS/MS. **C.** Levels of individual PA species. The data in B are shown here as the amounts of individual molecular species of PA (number of total carbons : number of double bonds). LHY/CCA1-binding PA species are highlighted in red. For B and C, values represent mean ± S.D. Asterisks denote significant difference from WT as determined by Student’s t-test (*p<0.05; **p<0.01; n=4 biological replicates). DW, dry weight of plant tissue.

### Molecular species of PA levels in the clock-altered PLD mutants

Since PA, the catalytic product of PLD, is an important regulator for the plant circadian clock, we analyzed PA contents in the 8 selected T3 plants to determine if and how they had changed. Total PA level in the mutants was highly consistent with the protein level of *PLDα1*; compared to WT, the total PA content was significantly reduced in *PLDα1*-mutated plants (20-7, 20-16, 25-12, and 25-14), whereas it remained essentially unchanged in the mutant plants without *PLDα1* mutated (7-11, 7-13, 16-12, and 16-13) (Figure 6B). Also worth noting is that the heterozygous *PLDα1* mutant 25-14 displayed about 3-fold higher total PA level than the homozygous *PLDα1* mutants, 20-7, 20-16, and 25-12 (Figure 6B). These data support that PLDα1 is the major contributor to PLD-catalyzed PA formation. Still, it was puzzling that the *PLDα1*-unmutated plants displayed the altered clock phenotype despite their comparable PA levels of WT. This prompted us to quantify individual molecular species of PA. Among 12 different PA species detected, levels of at least one of the two previously identified to associate with LHY and CCA1 (32:0 PA and 34:1 PA; Kim et al., 2019) were significantly decreased in all the mutants tested (Figure 6C). The *PLDα1* mutants also displayed a decrease in some other PA species, such as 34:2, 34:3, 36:2, 36:3, 36:4 and 36:5 PAs, but no other species was changed in the *PLDα1*-unmutated plants (Figure 6C). The data here suggest that specific PA species (*i.e.,* LHY/CCA1-binding PAs) generated by multiple PLDs, not limited to α1, are responsible for plant clock regulation.

### Changes in other membrane phospholipid species levels in the clock-altered PLD mutants

PLDs hydrolyze various membrane phospholipids into PA, with some displaying different substrate preferences for specific phospholipid classes (Yao et al., 2024). For example, PLDα1 preferentially hydrolyzes PC, whereas α3, γ1, δ, and ε use PE as a preferred substrate (Yao et al., 2024). Thus, we measured the levels of phospholipids to determine the lipid substrates that the PLD mutations might have accumulated at the expense of PA production in the 8 selected T3 plants. The reduced levels of total PA in *PLDα1* mutants were consistent with a significant accumulation of the major membrane phospholipids PC, PE, and phosphatidylglycerol (PG) in the same plants (Figure 7A). Noticeably, among all the *PLD* mutants analyzed, the mutant 20-7 showed an increase in phosphatidylserine (PS) content in addition to the major lipids (Figure 7A), possibly because it contains an additional *PLDδ* mutation and/or its unique combination of *PLDα1*, *δ*, and *ζ2* mutations (Figure 5B). By comparison, the other *PLD* mutants with unmutated *PLDα1* showed no change in the total content of phospholipid classes detected, with only 7-11 showing a slight increase in total PG level compared to WT (Figure 7A). However, lipid species analysis showed that the other *PLD* mutants were increased in the 32:0 and/or 34:1 PE and/or PG species, but not the 32:0 and 34:1 PC species. In contrast, the mutants containing mutated *PLDα1* all displayed increases in 32:0 PC, and they were increased in 34:1 PC species except for the heterogenous *PLDα1* 25-14 mutant (Figures 7B and S2; PE, PG, and PS species provided in Figures S3-5). The differential increases in different types and molecular species of phospholipids provide insights into potential substrates of the different PLDs involved in clock alterations. The differential accumulation of PC between *PLDα1*-mutated and unmutated plants suggests that PLDα1 hydrolyzes PC, as well as PE and PG, to generate 32:0 and 34:1 PA but the other PLDs hydrolyze 32:0 and 34:1 PE and PG species, but not PCs, to generate corresponding PA species.

**Figure 7.**
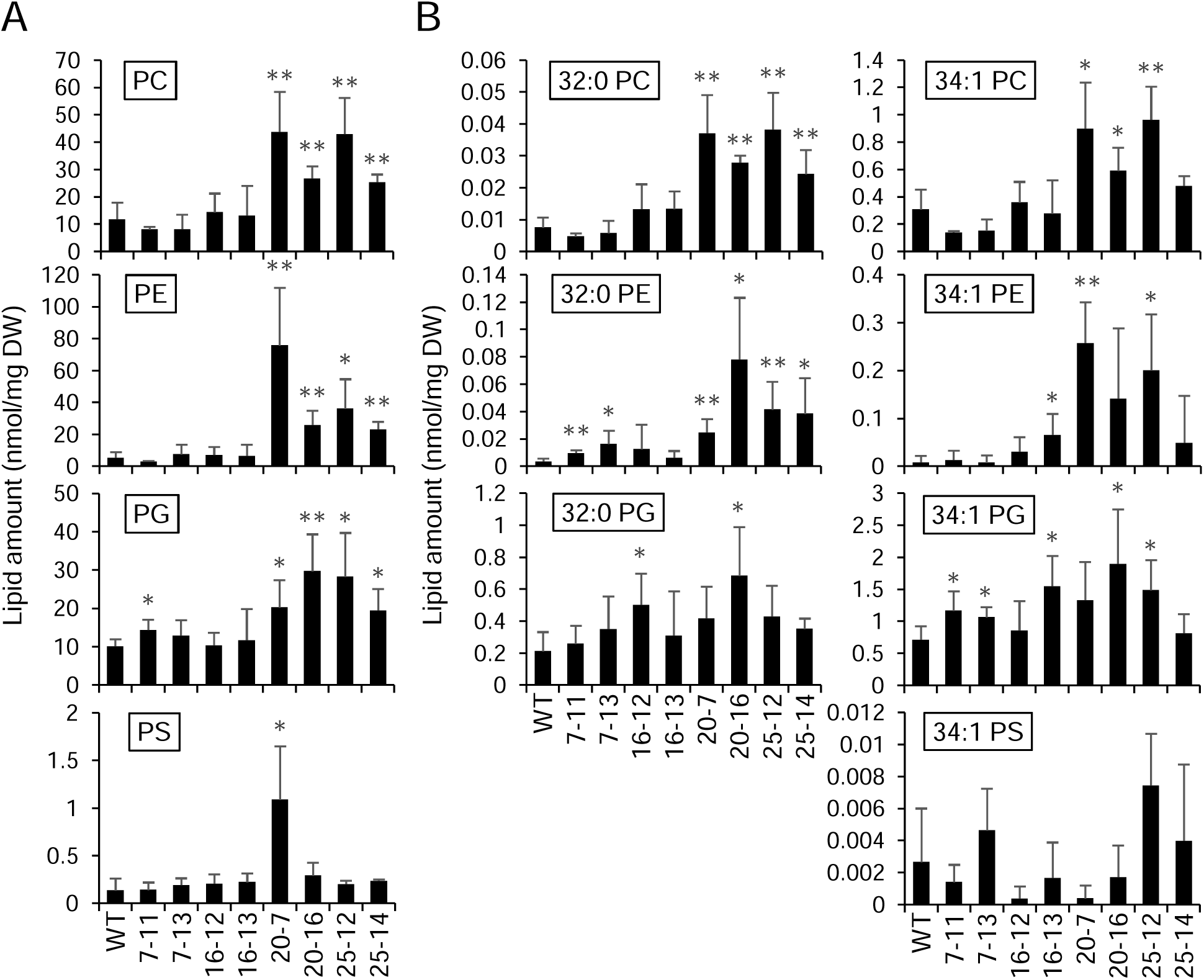
Phospholipid levels in the selected T3 mutants. Total lipids were extracted from the selected T3 mutants, and membrane phospholipids were separated and quantified by ESI-MS/MS. Each class of phospholipids is shown here as the amounts of total (A) and individual molecular species (32:0 and 34:1 (B); all other species provided in Supplemental figures 2-5). Note that 32:0 PS was not detectable. Values represent mean ± S.D. Asterisks denote significant difference from WT as determined by Student’s t-test (*p<0.05; **p<0.01; n=4 biological replicates). DW, dry weight of plant tissue.

## Discussion

Clock misalignments and mutations in clock genes lead to altered metabolism, whereas metabolic activities further affect the clock functions (Kim et al., 2019; Petrenko et al., 2023). The interplay between the clock and metabolism is expected to strengthen the function of both processes in organisms. Increasing evidence supports the clock regulation of plant lipid metabolism, including the recent findings that the core Arabidopsis clock morning regulators LHY/CCA1 enhance fatty acid biosynthesis and lipid accumulation (Kim et al., 2019, 2023). In contrast, the evening regulator TOC1 decreases lipid production (Makni et al., 2024). However, how lipid metabolism affects the molecular clock remains elusive. The results of the present study fill key knowledge gaps by identifying specific PLDs involved in modulating clock functions and provide valuable insights into the involvement of phospholipid species in the regulation of the clock function.

The results were made possible by generating a *PLD* mutant library using the multiplex CRISPR-Cas9 genome editing system. Our previous analysis of available single *PLD* null mutants and *pldα1pldδ* failed to identify PLDs involved in clock alterations (Kim et al., 2019). The potential compensation effects among 12 PLDs necessitate a more inclusive and unbiased approach to determine the clock-affecting PLDs. Although CRISPR-mediated genome editing provides a technical breakthrough for rapid and efficient gene alteration, its multiplex versions have been used only for simultaneous mutation of all desired genes in plants, mainly to decipher the function of a multigene family or metabolic pathway (Razzaq et al., 2019; Abdelrahman et al., 2021). A few mutant ‘libraries’ have been created using the CRISPR system, in tomato and rice, for example, but they were generated by polyclonal transformation of plants with a genomic-scale mixture of sgRNA constructs (Jacobs et al., 2017; Lu et al., 2017; Meng et al., 2017; Van Huffel et al., 2022). The polyclonal method could be used, but no attempts have been made to produce a plant library containing random combinations of mutations in a subset of or all members of a gene family. Construction of such a library could also be enabled by using a sgRNA with reduced specificity to target homologous genes, by which randomization may be challenging (Peng et al., 2015; Hyams et al., 2018). Recently, a multiplex CRISPR/Cas9 approach was developed for polycistronic assembly of sgRNA/Cas9 constructs for up to 32 sgRNAs (Stuttmann et al., 2021). Based on their finding that its editing efficiency was low in Arabidopsis, compared to tobacco plants, and limited with increasing numbers of sgRNAs, here we adapted the same vector system for one-shot generation of an Arabidopsis library mutated in various combinations of *PLD*s.

Our PCR and DNA sequencing analyses revealed that 11 of 12 *PLD*s were effectively edited by targeting two distinct sites, all heritable as demonstrated in the non-transgenic T2 plants. The editing of *PLDβ2* was undetected probably due to low on-target efficiency, compromised integrity (e.g. secondary structure), and/or low expression level of its sgRNAs (Javaid and Choi, 2021). Loss-of-function mutations of *PLDβ2* are unlikely to cause lethality because a knockdown mutant with a 77% reduction of its transcript showed no visible phenotypic difference from WT (Premkumar et al., 2019). More importantly, at least one across a wide spread of 11 *PLD*s was mutated in all T2 plants tested, suggesting that all 11 *PLD*s were subject to germ-line mutation. The T2 plants were only partially genotyped because the gel-based analysis was limited to detecting an insertion or deletion large enough to be resolved by electrophoretic separation of the PCR products. Despite the limitation, 1-3 out of 10 T2 plants (up to 30%) were found to bear mutations for each *PLD*. The editing frequency should be increased when analyzed by sequencing or other genotyping techniques, such as T7 endonuclease I, Surveyor nuclease, or PAGE-based genotyping assays (Yun et al., 2022). Nonetheless, we proceeded with screening our T2 plants to select clock-altered ones based on the effectiveness, heritability, and randomness of editing in *PLD*s.

Several plant circadian markers have been used to measure clock function, such as the rhythmic patterns in growth or organ movement (Tindall et al., 2015; Dakhiya et al., 2017). However, they require consistent positioning of individual plants and precise tracking of their organ positions on the time-course images, so they are not suited for screening large numbers of plants with a high time series resolution. One of the most widely used and reliable high-throughput methods is to monitor the activity of clock-controlled luciferase reporter, especially for plant species capable of stable transformation (Millar et al., 1992). Using this technique, here we measured the rhythmic patterns of the clock genes *CCA1* and *TOC1* expression in the T2 plants. The expression of *CCA1*/*LHY* and *TOC1* diurnally oscillates in an anti-phasic manner by the mutual repression mechanism (Alabadı et al., 2001; Gendron et al., 2012). Thus, clock dysfunction is often associated with an altered rhythmic expression of the clock genes mis-synchronized with the 24-hour periodic Earth’s rotation. Some of our T2 plants displayed a shorter *CCA1*/*TOC1* expression period, suggesting that they have the clock misaligned with the environmental rhythm.

Genotyping the progenies of the selected clock-altered plants reveals that at least 2 or 3 PLDs are required to alter the clock phenotype. This result is highly consistent with our previous observations that no clock-altering phenotype occurred in single T-DNA mutants of individual *PLD*s (Kim et al., 2019). Also, the *PLDζ2* mutation in addition to *α1* and *δ* in one of the plants (20-7 in Figure 5B) explains why LHY/CCA1-binding to *TOC1* promoter was not altered in *pldα1pldδ* (Kim et al., 2019). Together with these previous findings, our data here also indicate that for the PA regulation of the clock ‘specific’ PLDs (possibly the six mutated ones) are required, rather than just as many of ‘any’ PLDs as possible even though all PLDs produce PA, because no altered clock phenotype was observed when the two major PLDs (α1 and δ) were deficient. The importance of specific PLDs, not just the number of PLDs, is also supported by our phenotypic data that the T3 plants, which should have fewer *PLD*s mutated than their parental lines, still showed the altered clock phenotype.

The high frequency of mutations in *PLDα1*, *α3,* and *γ1* in the selected T3 plants is intriguing. Among the 12 PLDs in Arabidopsis, PLDα1 is the most abundant, highly ubiquitous, and functionally versatile isoform involved in responses to a variety of stimuli (Hong et al., 2016). *PLDα3* mutation is of interest because mass spectrometric lipid profiling in *pldα3* showed a dramatic decrease (∼90%) in the level of one of LHY/CCA1-binding PA species, 34:1-PA (Hong et al., 2008a). Also, PLDγ was found to be associated with the nuclear membrane, suggesting its potential role for PA formation in the nucleus where the clock transcription factors work (Fan et al., 1999). Thus, the three PLDs might significantly contribute to the clock function. Despite identifying the specific *PLD*s mutated, potential off-target effects should not be ruled out. Whole genome sequencing of Arabidopsis and rice has revealed that CRISPR/Cas9 system is highly specific in plants, unlike in human cells (Feng et al., 2014; Zhang et al., 2014; Wolt et al., 2016). In addition, deep sequencing of a total of 178 off-target sites showed that multiplex targeting in Arabidopsis resulted in no detectable off-target mutations (Peterson et al., 2016). Also, it is worth noting that the altered *CCA1* expression observed in T3 plants with heterozygous mutations could be caused by the negative dominant effect of the mutated PLDs and/or protein dosage effect, as evidenced for PLDα in plant 25-14. The biochemical and genetic basis for the clock output altered by the heterozygous, frame-shift mutations is under investigation.

The comparative lipidomic analysis of those clock-altering mutants and WT provides further insights into the lipid regulation of the circadian clock and PLD functions. It is not the level of total cellular PA, but the change in specific PA species associated with clock alterations. We showed that the LHY/CCA1 interactions with PA were species-specific, binding 16C-containing PA but not 18:1/18:1 PA (Kim et al., 2019). Even though the clock-altered *PLD* mutants without mutated *PLDα1* had comparable total PA levels as WT, all the clock-altered PLD mutants were decreased in the 16C-containing 32:0 and 34:1 PA. In addition, those mutants displayed corresponding increases in 32:0- and 34:1-containing PE, PG, and/or PC, but the increase differs among the mutants, indicating the potential substrate lipids for mutated PLDs. The clock-altered *PLD* mutants without mutated *PLDα1* were increased in 32:0- and 34:1-containing PE and PG, but not PC, species, whereas the *PLDα1*-mutated plants were all increased in 32:0- and 34:1 PC, as well other species. While *in vitro*, all PLDs can hydrolyze various membrane phospholipids into PA, it is highly challenging to decipher their lipid uses in plants. Limited substrate preference analyses suggest that PLDα1 preferentially hydrolyzes PC whereas α3, γ1, δ, and ε PE are preferred substrates (Yao et al., 2024). The present lipidomic analysis is mainly consistent with the PC preference by PLDα1, whereas the other *PLD* mutant combinations primarily hydrolyze PE and PG to generate the LHY/CCA1-interacting PA species.

Reciprocally, the circadian clock modulates lipid metabolism by regulating the expression of lipid metabolic genes, as evidenced by circadian profiles of the genes and oscillatory accumulation of some lipid metabolites, including PA, in plants (Maatta et al., 2012; Hsiao et al., 2014; Nakamura et al., 2014 a and b; Kim et al., 2019). Recently, we reported that LHY and CCA1 enhanced the expression of *KASIII* that encodes β-ketoacyl-ACP synthase III by directly binding to its promoter, and this binding was inhibited by PA (Kim et al., 2023). KASIII catalyzes the first condensation step in the biosynthetic pathway of fatty acids incorporated into myriad lipid species. The PA inhibition of LHY/CCA1-binding to the *KASIII* promoter suggests the role of PA as a retrograde modulator for the clock regulation of lipid biosynthesis. Together with the PA effects on circadian rhythms, this indicates that the PA-LHY/CCA1 interaction may serve as a cellular conduit to transmit signals between the clock and lipid metabolism in plants. Given the highly dynamic and complex nature of PA metabolism, discovering what controls the cellular levels of PA associated with the clock will help unveil how the two important processes communicate to affect plant performance.

## Materials and methods

### Cloning of sgRNA/Cas9 vector constructs

We used zCas9i cloning kit: MoClo-compatible CRISPR/Cas9 cloning kit with intronized Cas9 (AddGene #1000000171). The sequences of all oligonucleotides and PCR primers used in this study are provided in Table S1. Two target sequences for each *PLD* (23 bps including a PAM sequence) were selected by CHOPCHOP (https://chopchop.cbu.uib.no). For each of them two complementary oligonucleotides for sgRNA were synthesized to have 5’ (5’-ATTG-3’) and 3’ (5’-AAAC-3’) overhangs to be ligated with *BpiI* (*BbsI*)-cut shuttle vectors. They were hybridized by incubating at 95°C for 5 min and slowly cooling down to room temperature. Each step of the sequential cloning was performed by Golden Gate cloning with type II restriction enzymes and T4 DNA ligase. It was in a single tube reaction by thermal cycling (20-30 cycles of 37 °C for 2 min and 16 °C for 5 min) followed by incubation at 50 °C for 10 min and then 80 °C for 10 min. For shuttle vector cloning, 0.1 pmol of the hybridized oligonucleotides were cut by *BpiI* (or *BbsI*) and ligated with 0.2 μg of shuttle vectors (pDGE332, 333, 335, 336, 495, 496, 341, and 342) in a final volume of 10 μl. 0.5 μl of the reaction product was transformed into *E. coli* DH5α, and all transformed cells were grown together in 3 ml LB with carbenicillin at 37 °C overnight for polyclonal plasmid preparation. For multi-multi vector cloning, 1 μl of the 8 shuttle vector constructs were cut by *BsaI* and ligated with 0.2 μg of multi-multi vectors (pDGE503, 504, and 505) in a final volume of 20 μl. 2 μl of the reaction product was transformed into *E. coli* DH5α, and kanamycin-resistant clones were obtained. Plasmids were extracted from the *E. coli* clones selected to have all 8 sgRNAs. For recipient vector cloning, 2 μl of the three multi-multi vector constructs were cut by *BpiI* and *BsaI* and ligated with 0.3 μg of recipient vector (pDGE668) and 30 ng of end-linker vector (pDGE509) in a final volume of 20 μl. *E. coli* transformation was done to select spectinomycin-resistant clones. Plasmids were extracted from the *E. coli* clones with all 24 sgRNAs and used to transform *Arabidopsis thaliana* (Col-0) via Agrobacterium-mediated transformation by floral dipping.

### Confirmation of cloning at each step

For confirmation of each of the 24 shuttle vector constructs, plasmid was extracted using High-Speed Plasmid Mini Kit (IBI Scientific) polyclonally from the 3 ml culture of pooled *E. coli* transformants, cut with 1 unit of *BamHI* (Ficher) at 37 °C for 15 min, and analyzed in 1.5% agarose gel. To confirm each of the 3 multi-multi vector constructs, 5 randomly selected *E. coli* colonies were grown individually in 3 ml LB at 37 °C overnight. Plasmid was extracted from each of them, cut with 1 unit of *BpiI* (NEB) at 37 °C for 1 hour, and analyzed in 1.5% agarose gel. For the recipient vector construct, plasmids from 4 randomly selected *E. coli* clones were cut with 1 unit of *HindII* (Fisher) at 37 °C for 15 min and analyzed in 0.8% agarose gel. All agarose gels were stained with ethidium bromide to visualize DNA under UV light.

### PCR analysis and sequencing of genomic DNA

Genomic DNA was extracted from a small piece of Arabidopsis leaf using Extract-N-Amp extraction kit (Sigma-Aldrich) and used as a template for PCR. Reaction mixture was set in a final volume of 20 μl as 0.2 μM dNTP mix, 50 μM each primer, 0.5 μl template, 1 unit of Taq DNA polymerase (GenScript). PCR was conducted at 94 °C for 3 min for denaturation followed by 35-40 cycles of 94 °C for 30 sec, 50-55 °C for 45 sec and 72 °C for 60 sec, with final extension at 72 °C for 7 min. PCR products were analyzed in 1% agarose gel stained with ethidium bromide. For DNA sequencing, the PCR products were purified using Gel/PCR DNA Fragments Extraction Kit (IBI Scientific) and sent with a 10 μM primer (forward and/or reverse depending on the chromatogram quality) for a commercial Sanger sequencing service (Eurofins Genomics). Sequencing results were processed and analyzed using Chromas software (v 2.6.6).

### Selection of non-transgenic seeds

T2 seeds were harvested and approximately 200-300 seeds were evenly spread on a white paper. They were mounted and observed in a fluorescence microscope with RFP filter (Eclipse E800, Nikon). The seeds without fluorescence were carefully picked up using the tip of a wet tooth pick and transferred to water in a microtube for sterilization as described below. This process was repeated until a sufficient amount of non-transgenic seeds was obtained.

### Plant materials and growth conditions

All Arabidopsis plants used in this study were Col-0 ecotype of *Arabidopsis thaliana*. Transgenic plants with *CCA1::LUC* or *TOC1::LUC* were previously generated (Kim et al., 2019). Seeds were surface-sterilized with 70 % (v/v) ethanol and then with 20 % (v/v) bleach, washed with water 4 times, and sown on 1/2 strength of Murashige and Skoog (MS) media supplemented with 1 % (w/v) sucrose and 0.8 % (w/v) agar. After stratification at 4 °C for 2 days in the dark, plants were germinated and grown in a growth chamber (Percival, CU22L2) maintained at 22 °C and a relative humidity of 60% under light cycles of 12-h light/12-h dark with a photosynthetic photon flux density of 120-150 μmol/m^2^/sec.

### Time-course measurement of luciferase activity

The *CCA1::LUC* and *TOC1::LUC* constructs were previously described (Alabadí et al., 2001). Sterilized seeds were sown on 1/2 MS plate (10×10 cm) supplemented with 1% sucrose in lines with intervals of at least 1.5 cm. The plates were sealed with paper tapes, not Parafilm, to promote gas exchange and prevent condensation on the lid. The seeds were germinated and grown for entrainment in 12-h light/12-h dark cycles at 22 °C for 7 days. Under aseptic conditions, they were sprayed evenly with 5 μM luciferin dissolved in 0.01% Triton-X100 and transferred to a growth chamber equipped with PIXIS 1024B CCD camera (Princeton Instruments) and LED lights (70 μmol/m^2^/s, wavelengths 400, 430, 450, 530, 630, and 660 at intensity 350; Heliospectra LED lights). Plant images were taken every 60 min after a 180-sec delay for 4 min with 1×1 binning using µManager software (Edelstein et al., 2010, 2014), starting at Zeitgeber time (ZT) 0 hour for *CCA1::LUC* and 12 hour for *TOC1::LUC* to synchronize the phase of their expression. The luminescence intensity from each seedling on each image was measured with ImageJ (1.53e) using the ROI manager. Circadian parameters were estimated using BioDare2 (biodare2.ed.ac.uk) with linear detrend and the fast fourier transform non-linear least squares (FFT-NLLS) method (Zieliński et al., 2021).

### Protein analysis by immunoblotting

Plant tissues were ground with liquid nitrogen and mixed with protein extraction buffer (50 mM Tris pH7.3, 50 mM NaCl, 5 % glycerol). The tissue homogenate was filtered through 4 layers of Miracloth® (Calbiochem) and centrifuged at 12,000 xg for 10 min at 4 °C. The supernatant was mixed with SDS-PAGE loading buffer, boiled for 5 min, and resolved in 10 % (v/v) polyacrylamide gel at 100 V for ∼1 hour. Proteins were electrophoretically transferred onto nitrocellulose membrane using Semidry Trans-Blot apparatus (Bio-Rad) at 20 V for 20 min. The membrane was blocked in tris-buffered saline with 0.1 % (v/v) Tween-20 (TBST) containing 5 % (w/v) nonfat milk for 1 hour, followed by washing three times with TBST buffer. The membrane was incubated with primary antibodies (anti-PLDα or anti-GAPC) for 1 hour. After washing three times with TBST buffer, the membrane was incubated for 1 hour with a secondary antibody (anti-rabbit) conjugated with horseradish peroxidase. Proteins were visualized using SuperSignal^TM^ West Pico PLUS chemiluminescent substrate (Thermo Scientific) and imaged by iBright 1500 imager (Invitrogen).

### Lipid analysis by mass spectrometry

Leaves from ∼4-week-old plants were incubated with 3 mL isopropanol containing 0.01 % (w/v) butylated hydroxytoluene (BHT) at 75 °C for 15 min to prevent lipoxidation and lipolysis. Total lipids were extracted from the tissues by agitating with 1 mL chloroform and 0.6 mL water for 1 hour. Lipids were further extracted three times with 3 mL of a 2:1 (v/v) mixture of chloroform and methanol containing 0.01% (w/v) BHT, with collecting the organic phase into a fresh tube each time. All lipid extracts were combined and washed twice by mixing with 1 mL of 1 M KCl (water for the second wash) and discarding the upper aqueous phase after a brief centrifugation. The lower organic phase was dried under nitrogen gas and re-dissolved in 1 mL chloroform. The resulting lipids were applied to electrospray ionization (ESI)-MS/MS (API-4000, SCIEX, Framingham, MA) detection system with a mixture of internal lipid standards and solution B (95 % (v/v) methanol, 14.3 mM ammonium acetate). After lipid extraction, the remaining tissues were air-dried and measured for weight. Data were processed and quantified by Analyst software (v1.5.1). Lipid contents were calculated as molar amounts per tissue dry weight.

## Acknowledgements

Research reported in this article was supported by the National Institute of General Medical Sciences of the National Institutes of Health under award number R01GM141374. The authors declare no conflict of interest.

## Author contributions

S.K. designed and performed all experiments, collected and analyzed all data, and wrote the manuscript. D.N. discussed the results and edited the manuscript. X.W. proposed and supervised the study, discussed the results, and edited the manuscript.

## Data availability

The data that support the findings of this study are available from the corresponding author upon reasonable request.

## References

Abdelrahman, M., Wei, Z., Rohila, J.S. and Zhao, K., 2021. Multiplex genome-editing technologies for revolutionizing plant biology and crop improvement. Frontiers in Plant Science, 12, p.721203.

Alabadı, D., Oyama, T., Yanovsky, M.J., Harmon, F.G., Más, P. and Kay, S.A., 2001. Reciprocal regulation between TOC1 and LHY/CCA1 within the Arabidopsis circadian clock. Science, 293(5531), pp.880–883.

Ali, U., Lu, S., Fadlalla, T., Iqbal, S., Yue, H., Yang, B., Hong, Y., Wang, X. and Guo, L., 2022. The functions of phospholipases and their hydrolysis products in plant growth, development and stress responses. Progress in Lipid Research, 86, p.101158.

Buckley, C.R., Li, X., Martí, M.C. and Haydon, M.J., 2023. A bittersweet symphony: Metabolic signals in the circadian system. Current Opinion in Plant Biology, 73, p.102333.

Budai, Z., Balogh, L. and Sarang, Z., 2019. Short-term high-fat meal intake alters the expression of circadian clock-, inflammation-, and oxidative stress-related genes in human skeletal muscle. International Journal of Food Sciences and Nutrition, 70(6), pp.749–758.

Dakhiya, Y., Hussien, D., Fridman, E., Kiflawi, M. and Green, R., 2017. Correlations between circadian rhythms and growth in challenging environments. Plant Physiology, 173(3), pp.1724–1734.

Deepika, D. and Singh, A., 2022. Plant phospholipase D: novel structure, regulatory mechanism, and multifaceted functions with biotechnological application. Critical Reviews in Biotechnology, 42(1), pp.106–124.

Delezie, J. and Challet, E., 2011. Interactions between metabolism and circadian clocks: reciprocal disturbances. Annals of the New York Academy of Sciences, 1243(1), pp.30–46.

Edelstein, A., Amodaj, N., Hoover, K., Vale, R. and Stuurman, N., 2010. Computer control of microscopes using µManager. Current Protocols in Molecular Biology, 92(1), pp.14–20.

Edelstein, A.D., Tsuchida, M.A., Amodaj, N., Pinkard, H., Vale, R.D. and Stuurman, N., 2014. Advanced methods of microscope control using μManager software. Journal of Biological Methods, 1(2).

Fan, L., Zheng, S., Cui, D. and Wang, X., 1999. Subcellular distribution and tissue expression of phospholipase Dα, Dβ, and Dγ in Arabidopsis. Plant Physiology, 119(4), pp.1371–1378.

Feng, Z., Mao, Y., Xu, N., Zhang, B., Wei, P., Yang, D.L., Wang, Z., Zhang, Z., Zheng, R., Yang, L. and Zeng, L., 2014. Multigeneration analysis reveals the inheritance, specificity, and patterns of CRISPR/Cas-induced gene modifications in Arabidopsis. Proceedings of the National Academy of Sciences, 111(12), pp.4632–4637.

Gendron, J.M., Pruneda-Paz, J.L., Doherty, C.J., Gross, A.M., Kang, S.E. and Kay, S.A., 2012. Arabidopsis circadian clock protein, TOC1, is a DNA-binding transcription factor. Proceedings of the National Academy of Sciences, 109(8), pp.3167–3172.

Grosjean, E., Simonneaux, V. and Challet, E., 2023. Reciprocal interactions between circadian clocks, food intake, and energy metabolism. Biology, 12(4), p.539.

Grützner, R., Martin, P., Horn, C., Mortensen, S., Cram, E.J., Lee-Parsons, C.W., Stuttmann, J. and Marillonnet, S., 2021. High-efficiency genome editing in plants mediated by a Cas9 gene containing multiple introns. Plant Communications, 2(2).

Hong, Y., Pan, X., Welti, R. and Wang, X., 2008(a). Phospholipase Dα3 is involved in the hyperosmotic response in Arabidopsis. The Plant Cell, 20(3), pp.803–816.

Hong, Y., Zheng, S. and Wang, X., 2008(b). Dual functions of phospholipase Dα1 in plant response to drought. Molecular Plant, 1(2), pp.262–269.

Hong, Y., Zhao, J., Guo, L., Kim, S.C., Deng, X., Wang, G., Zhang, G., Li, M. and Wang, X., 2016. Plant phospholipases D and C and their diverse functions in stress responses. Progress in Lipid Research, 62, pp.55–74.

Hsiao, A.S., Haslam, R.P., Michaelson, L.V., Liao, P., Napier, J.A. and Chye, M.L., 2014. Gene expression in plant lipid metabolism in Arabidopsis seedlings. PLoS One, 9(9), p.e107372.

Hyams, G., Abadi, S., Lahav, S., Avni, A., Halperin, E., Shani, E. and Mayrose, I., 2018. CRISPys: optimal sgRNA design for editing multiple members of a gene family using the CRISPR system. Journal of Molecular Biology, 430(15), pp.2184–2195.

Jacobs, T.B., Zhang, N., Patel, D. and Martin, G.B., 2017. Generation of a collection of mutant tomato lines using pooled CRISPR libraries. Plant Physiology, 174(4), pp.2023–2037.

Javaid, N. and Choi, S., 2021. CRISPR/Cas system and factors affecting its precision and efficiency. Frontiers in Cell and Developmental Biology, 9, p.761709.

Kim, S.C., Nusinow, D.A., Sorkin, M.L., Pruneda-Paz, J. and Wang, X., 2019. Interaction and regulation between lipid mediator phosphatidic acid and circadian clock regulators. The Plant Cell, 31(2), pp.399–416.

Kim, S.C. and Wang, X., 2020. Phosphatidic acid: an emerging versatile class of cellular mediators. Essays in Biochemistry, 64(3), pp.533–546.

Kim, S.C., Edgeworth, K.N., Nusinow, D.A. and Wang, X., 2023. Circadian clock factors regulate the first condensation reaction of fatty acid synthesis in Arabidopsis. Cell Reports, 42(12).

Kim, S.C., Yao, S., Zhang, Q. and Wang, X., 2022. Phospholipase Dδ and phosphatidic acid mediate heat induced nuclear localization of glyceraldehydeL3Lphosphate dehydrogenase in Arabidopsis. The Plant Journal, 112(3), pp.786–799.

Kocourková, D., Kroumanová, K., Podmanická, T., Daněk, M. and Martinec, J., 2021. Phospholipase Dα1 acts as a negative regulator of high Mg^2+^-induced leaf senescence in Arabidopsis. Frontiers in Plant Science, 12, p.770794.

Kolesnikov, Y., Kretynin, S., Bukhonska, Y., Pokotylo, I., Ruelland, E., Martinec, J. and Kravets, V., 2022. Phosphatidic acid in plant hormonal signaling: from target proteins to membrane conformations. International Journal of Molecular Sciences, 23(6), p.3227.

Liu, Q., Zhou, Y., Li, H., Liu, R., Wang, W., Wu, W., Yang, N. and Wang, S., 2021. Osmotic stress-triggered stomatal closure requires Phospholipase Dδ and hydrogen sulfide in Arabidopsis thaliana. Biochemical and Biophysical Research Communications, 534, pp.914–920.

Lu, Y., Ye, X., Guo, R., Huang, J., Wang, W., Tang, J., Tan, L., Zhu, J.K., Chu, C. and Qian, Y., 2017. Genome-wide targeted mutagenesis in rice using the CRISPR/Cas9 system. Molecular Plant, 10(9), pp.1242–1245.

Maatta, S., Scheu, B., Roth, M.R., Tamura, P., Li, M., Williams, T.D., Wang, X. and Welti, R., 2012. Levels of Arabidopsis thaliana leaf phosphatidic acids, phosphatidylserines, and most trienoate-containing polar lipid molecular species increase during the dark period of the diurnal cycle. Frontiers in Plant Science, 3, p.49.

Makni S, Acket S, Guenin S, Afensiss S, Guellier A, Martins-Noguerol R, Moreno-Perez AJ, Thomasset B, Martinez-Force E, Gutierrez L, Ruelland E, Troncoso-Ponce A. Arabidopsis seeds altered in the circadian clock protein TOC1 are characterized by higher level of linolenic acid. Plant Sci. 2024 Apr 9;344:112087. doi: 10.1016/j.plantsci.2024.112087. Epub ahead of print. PMID: 38599247.

McClung, C.R., 2019. The plant circadian oscillator. Biology, 8(1), p.14.

Meng, X., Yu, H., Zhang, Y., Zhuang, F., Song, X., Gao, S., Gao, C. and Li, J., 2017. Construction of a genome-wide mutant library in rice using CRISPR/Cas9. Molecular Plant, 10(9), pp.1238–1241.

Millar, A.J., Short, S.R., Chua, N.H. and Kay, S.A., 1992. A novel circadian phenotype based on firefly luciferase expression in transgenic plants. The Plant Cell, 4(9), pp.1075–1087.

Munnik, T., Arisz, S.A., De Vrije, T. and Musgrave, A., 1995. G protein activation stimulates phospholipase D signaling in plants. The Plant Cell, 7(12), pp.2197–2210.

Nakamura, Y., Andrés, F., Kanehara, K., Liu, Y.C., Coupland, G. and Dörmann, P., 2014(a). Diurnal and circadian expression profiles of glycerolipid biosynthetic genes in Arabidopsis. Plant Signaling & Behavior, 9(9), p.e29715.

Nakamura, Y., Andrés, F., Kanehara, K., Liu, Y.C., Dörmann, P. and Coupland, G., 2014(b). Arabidopsis florigen FT binds to diurnally oscillating phospholipids that accelerate flowering. Nature Communications, 5(1), pp.1–7.

Ordon, J., Gantner, J., Kemna, J., Schwalgun, L., Reschke, M., Streubel, J., Boch, J. and Stuttmann, J., 2017. Generation of chromosomal deletions in dicotyledonous plants employing a user friendly genome editing toolkit. The Plant Journal, 89(1), pp.155–168.

Pan, X., Mota, S. and Zhang, B., 2020. Circadian clock regulation on lipid metabolism and metabolic diseases. Lipid Transfer in Lipoprotein Metabolism and Cardiovascular Disease, pp.53–66.

Peng, D., Kurup, S.P., Yao, P.Y., Minning, T.A. and Tarleton, R.L., 2015. CRISPR-Cas9-mediated single-gene and gene family disruption in Trypanosoma cruzi. MBio, 6(1), pp.10–1128.

Peterson, B.A., Haak, D.C., Nishimura, M.T., Teixeira, P.J., James, S.R., Dangl, J.L. and Nimchuk, Z.L., 2016. Genome-wide assessment of efficiency and specificity in CRISPR/Cas9 mediated multiple site targeting in Arabidopsis. PloS One, 11(9), p.e0162169.

Premkumar, A., Lindberg, S., Lager, I., Rasmussen, U. and Schulz, A., 2019. Arabidopsis PLDs with C2 domain function distinctively in hypoxia. Physiologia Plantarum, 167(1), pp.90–110.

Qin, C. and Wang, X., 2002. The Arabidopsis phospholipase D family. Characterization of a calcium-independent and phosphatidylcholine-selective PLDζ1 with distinct regulatory domains. Plant Physiology, 128(3), pp.1057–1068.

Qu, L., Chu, Y.J., Lin, W.H. and Xue, H.W., 2021. A secretory phospholipase D hydrolyzes phosphatidylcholine to suppress rice heading time. PLoS Genetics, 17(12), p.e1009905.

Razzaq, A., Saleem, F., Kanwal, M., Mustafa, G., Yousaf, S., Imran Arshad, H.M., Hameed, M.K., Khan, M.S. and Joyia, F.A., 2019. Modern trends in plant genome editing: an inclusive review of the CRISPR/Cas9 toolbox. International Journal of Molecular Sciences, 20(16), p.4045.

Song, P., Jia, Q., Chen, L., Jin, X., Xiao, X., Li, L., Chen, H., Qu, Y., Su, Y., Zhang, W. and Zhang, Q., 2020. Involvement of Arabidopsis phospholipase Dδ in regulation of ROS-mediated microtubule organization and stomatal movement upon heat shock. Journal of Experimental Botany, 71(20), pp.6555–6570.

Stuttmann, J., Barthel, K., Martin, P., Ordon, J., Erickson, J.L., Herr, R., Ferik, F., Kretschmer, C., Berner, T., Keilwagen, J. and Marillonnet, S., 2021. Highly efficient multiplex editing: one shot generation of 8× Nicotiana benthamiana and 12× Arabidopsis mutants. The Plant Journal, 106(1), pp.8–22.

Su, Y., Li, M., Guo, L. and Wang, X., 2018. Different effects of phospholipase Dζ2 and non specific phospholipase C4 on lipid remodeling and root hair growth in Arabidopsis response to phosphate deficiency. The Plant Journal, 94(2), pp.315–326.

Tindall, A.J., Waller, J., Greenwood, M., Gould, P.D., Hartwell, J. and Hall, A., 2015. A comparison of high-throughput techniques for assaying circadian rhythms in plants. Plant Methods, 11, pp.1–7.

Van Huffel, K., Stock, M., Ruttink, T. and De Baets, B., 2022. Covering the combinatorial design space of multiplex CRISPR/Cas experiments in plants. Frontiers in Plant Science, 13, p.907095.

Wolt, J.D., Wang, K., Sashital, D. and LawrenceLDill, C.J., 2016. Achieving plant CRISPR targeting that limits off target effects. The Plant Genome, 9(3), pp.plantgenome2016-05.

Yao, S., Peng, S. and Wang, X., 2022(a). Phospholipase Dε interacts with autophagy related protein 8 and promotes autophagy in Arabidopsis response to nitrogen deficiency. The Plant Journal, 109(6), pp.1519–1534.

Yao, S., Wang, G. and Wang, X., 2022(b). Effects of phospholipase dε overexpression on soybean response to nitrogen and nodulation. Frontiers in Plant Science, 13, p.852923.

Yun, J.Y., Yu, S.I., Bang, S.E., Kim, J.Y., Lee, S.H. and Lee, B.H., 2022. Identification of CRISPR-induced mutations in plants: with a focus on the next-generation sequencing assay. Journal of Plant Biology, 65(6), pp.435–443.

Zhang, G., Yang, J., Chen, X., Zhao, D., Zhou, X., Zhang, Y., Wang, X. and Zhao, J., 2021. Phospholipase DL and phosphatidic acid mediated phospholipid metabolism and signaling modulate symbiotic interaction and nodulation in soybean (Glycine max). The Plant Journal, 106(1), pp.142–158.

Zhang, H., Zhang, J., Wei, P., Zhang, B., Gou, F., Feng, Z., Mao, Y., Yang, L., Zhang, H., Xu, N. and Zhu, J.K., 2014. The CRISPR/C as9 system produces specific and homozygous targeted gene editing in rice in one generation. Plant Biotechnology Journal, 12(6), pp.797–807.

Zieliński, T., Hay, J. and Millar, A.J., 2021. Period estimation and rhythm detection in timeseries data using biodare2, the free, online, community resource. In Plant Circadian Networks: Methods and Protocols (pp. 15–32). New York, NY: Springer US.

