## Supplement figures for "Identification of phospholipase Ds and phospholipid species involved in circadian clock alterations using CRISPR/Cas9-based multiplex editing of Arabidopsis"

PLDα1

ATGGCGCAGCATCTGTTGCACGGGACTTTACATGCTACCATCTATGAAGTTGATGCCCTCCATGGTGGTGGTGTTAGGCA
AGGCTTCCTTGGCAAGGTATAATAAGATCCATTGTTATTGCATCATTTTTGCGCCAATCCTGACGTCTTATTTTGTAGTT
CAAATTAGTCTGTGGACTAATACACAGTTTGATCTTTATTTTTCTGATGAAACTTTCAGATTCTGGCAAATGTAGAAGAG
ACGATTGGTGTTGGTAAAGGAGAAACACAGTTGTATGCGACGATTGATCTGCAAAAAGCTAGAGTTGGGAGAACCAGGAA
GATCAAAAATGAACCTAAGAACCCAAAGTGGTATGAGTCGTTTCATATTTACTGTGCTCACTTGGCTTCTGATATCATCT
TCACTGTTAAAGATGATAATCCCATTGGAGCTACCCTTATCGGAAGAGCTTACATTCCTGTTGATCAAGTCATTAACCGGC
GAGGAAGTGGATCAGTGGGTTGAGATCTTGGATAATGACAGAAACCCTATTCAGGGAGGATCAAAGATTCATGTCAAGCT

PLDα2

ATGGAAGAGTGTTTGTTACATGGAAGATTGCACGCGACAATCTACGAAGTTGATCATCTCCATGCTGAAGGAGGTAGATC
AGGTTTCTTAGGATCGGTATTGTCTTTTGTCTCTCTCTCATTTATAAGTTGTTTAGAGATACAACAACAAAAGGTTTCCT
TCAATCTGTCTTTAATGTTCTGTTTTTAATCTTTCCAGATACTAGCAAATGTAGAAGAAACAATCGGTGTTGGCAAAGGA
GAAACTCAGCTTTACGCTACAATCGATCTTGAGAAAGCAAGAGTTGGAAGAACGAGGAAAATAACAAAGGAGCCAAAAAA
CCCAAAGTGGTTTGAGTCTTTTCATATTTACTGTGGTCACATGGCTAAACATGTTATTTTCACTGTTAAGGATGCTAATC
CGATTGGCGCGACGTTGATCGGAAGAGGTTATATTCCTGTGGAAGATATTTTGCATGGAGAAGAAGTTGATAGATGGGTT
GATATCTTAGATAATGAGAAAAACCCAATTGCTGGAGGATCAAAGATCCATGTGAAACTCCAATATTTCGGTGTTGAAAA
GGATAAAAACTGGAATCGTGGTATCAAAAGTGCTAAGTTTCCAGGAGTTCCTTACACATTCTTCTCTCAGAGGAGAGGCT

PLDα3

ATGACGGAGCAATTGCTGCTTCATGGAACTCTTGAAGTGAAGATTTATAGGATCGATAAGTTGCATCAACGTTCAAGATT
CAATTTATGCGGCAAGGTAATATTTAATTTCAACACTATCTACTAATAAGAGATATTAGACGAAGGTTACTTGTAAAATA
AGACTCAAAAATGATTCATAGGGGAACAAGGAGCCAACAGGTAAAAAGACTCAATCTCAAATCAAAAGACTAACGGACTC
ATGCACAAGTTTGTTTGGAGGACATCTATACGCAACTATTGACCTAGACAGGTCAAGAGTGGCTAGAACCATGATGAGAC
GTCATCCTAAATGGTTACAATCCTTCCACGTCTACACCGCACATTCAATCTCTAAAATCATATTCACAGTTAAAGAAGAT
GAACCGGTCAGTGCTAGTTTGATAGGGCGAGCTTATTTACCTGTAACAGAAGTCATCACCGGACAACCTATAGACCGATG
GCTCGATATCTTAGACGAGAACAGGAGACCAATCCAAGGAGGTTCCAAACTCCATGTACGTGTGAAGTTCACTCATGTGA
CACAGGACGTGAATTGGAACAAAGGAATAATATTACCTTCCTTCAATGGAGTTCCTAACGCTTATTTCAACCAAAGAGAA
GGTTGCAAAGTTACACTATACCAAGACGCTCACGTTCTTAACGAATACCCTGATGTTACCCTCACGGGAGGACAAGTTAT
ATACAAGCATCATCGGTGTTGGGAAGAGATTTTCGATGCGATATGGGAAGCAAAACATTTGATATACATCGCTGGTTGGT

PLDβ1

ATGGATAATCACGGTCCTCGTTATCCATACCCTTACGGTCAGTACCCATACCCTTACCCATACCCAGCTCCTTATAGACC
TCCCAGTTCAGAGCCATACCCACCTCCTCCAACCAATCAATACAGTGCTCCTTATTACCCTTACCCACCACCTCCATACG
CAACACCACCACCATATGCATCACCACCACCGCCTCATCAGCACACTTCGGGTTCGCATTCTGGACCATTAGACTATAGC
CACAACCCACAACCATCATCTCTCGCGGCTGCTCCTCCAGAGTATCACAGACATTCTTTTGACTACCAACCTTCTCCTTA
CCCTTATCAGCCCCAAGGTAACTTTGGTGCTTATGGTCCTCCGCCTCCTCACTATTCGTATCAAGAGCCAGCTCAATACC
CTCCACCTGAAACTAAACCGCAAGAGCCTCTGCCTCCTCCGCAGCAAACTCAAGGTTTTCAAGAATATCGTAGGCAAGAT
TGTCTCAGTACCGGGGGAACAGGTCATGATAATGTAAGCAATTCTGGATCTTCTTATCCTCCTGTGGATGAACTTCTAGG
TGGTCTGCATATTTCTACTAACCAACCAGGTCCCTCAGTTCCACAACTATCCTCCCTTCCTTCGAACTCTTGGCAAAGCC

PLDβ2

ATGGAGAATTATGGTTGGAATTACCCTTATTACCCTTACAGGCCTCCGCGGCCTAATCCTCCTTACCCAGCACCGCCTCA
TCACCATGGATCAATGTCTCATTCGGGTCCATTAGACCACCACCACCCTCCAATGAGTTACTATGCTTCTTTTGATTACC
AACACCAACCTCCTCCTCCTTACCCTCCAGTGAGTTACTATGCTTCGTTTTCAAGTCATTCAGATTTATCATACTCAGGC
AGGCTAGATTCGTCTGGTCATGGTTTTACATCTACCGCATCGCCGCATAGTCCCGGTATGCACATTGTGCCCTTTGGTAA
AGCCTCCTTGAAGGTCCTCTTGTTGCATGGTAATTTGGATATTTGGGTTTCTTGTGCTAATAATCTACCAAACTTAGATT
TGTTTCATAAGACATTGGGTGTTGTGTTTGGTGGCATGACCAATATGATCGAAGGCCAGTTGAGCAAGAAGATCACTAGT
GATCCTTACGTGTCCATATCAGTCGCTGGTGCTGTCATAGGAAGAACATATGTCATTAGCAACAGCGAAAATCCTGTCTG
GCAGCAGCATTTTTATGTCCCTGTAGCTCATCACGCTGCCGAGGTCCATTTTGTTGTTAAGGACAGTGATGCCGTCGGTT
CACAGCTTATAGGAATTGTTACAATCCCTGTTGAACAAATTTACTCAGGTGCAAGAATTGAAGGGACTTACTCCATTCGA

PLDγ1

ATGGCGTATCATCCGGCTTATACTGAGACTATGTCTATGGGAGGAGGGTCAAGCCATGGGGGCGGCCAGCAATACGTGCC
GTTTGCTACGAGTAGTGGCTCTTTGAGGGTCGAATTGTTACATGGTAACTTAGACATTTGGGTTAAGGAAGCTAAACATC
TTCCTAATATGGATGGTTTCCATAACAGACTTGGTGGGATGTTGTCTGGGTTAGGTCGGAAGAAAGTGGAAGGTGAGAAG
TCTTCTAAGATCACTAGTGATCCTTATGTTACTGTCTCTATCTCTGGTGCTGTGATTGGTAGAACTTTTGTTATTAGCAA
TAGTGAGAATCCTGTGTGGATGCAGCATTTTGATGTACCGGTTGCTCATAGTGCTGCTGAAGTACATTTTGTTGTGAAAG
ATAGTGATATTATTGGATCACAGATCATGGGAGCTGTTGGAATCCCAACGGAGCAGCTGTGTTCGGGGAATAGGATCGAA

PLDγ2

ATGTCAATGGGAGGAGGGTCAAACCACGAGTTTGGCCAGTGGCTTGACCAGCAACTCGTTCCGTTAGCTACGAGTAGTGG
CTCTTTGATGGTTGAATTGTTACATGGTAACTTAGACATTTGGGTTAAGGAAGCTAAACATCTTCCTAACATGATATGTT
ACCGTAACAAGCTTGTTGGTGGGATTTCGTTTTCTGAGTTAGGTCGGAGGATTCGTAAAGTGGATGGTGAGAAGTCTTCT
AAGTTCACAAGTGATCCTTATGTTACTGTCTCTATCTCTGGTGCTGTCATTGGTAGAACTTTTGTTATTAGCAATAGTGA
GAATCCTGTGTGGATGCAGCATTTCGATGTACCCGTTGCTCATAGTGCTGCTGAAGTACATTTTGTTGTGAAAGACAATG

PLDγ3

ATGGCGTATCATCCAGTTTATAACGAGACTATGTCTATGGGAGGAGGGTCAAGCAACGAGTTTGGCCAGTGGCTTGACAA
GCAACTCGTGCCGTTTGATACAAGTAGTGGCTCTTTGAGGGTTGAATTATTACATGGTAACTTAGACATTTGGGTTAAGG
AAGCTAAACATCTTCCTAACATGGATGGTTTCCATAACACCCTTGTTGGTGGGATGTTTTTTGGGTTAGGTCGGAGGAAT
CATAAAGTGGATGGTGAGAATTCTTCTAAGATCACAAGTGATCCTTATGTTACTGTCTCTATCTCTGGTGCTGTGATTGG
TAGAACTTTTGTTATTAGCAATAGTGAGAATCCTGTGTGGATGCAGCATTTTGATGTACCAGTTGCTCATAGTGCTGCGA
AAGTTCACTTTGTTGTGAAAGACAGTGATATTATTGGATCACAGATCATTGGAGCTGTTGAAATCCCAACCGAGCAGTTG
TGTTCCGGGAATAGAATCGAAGGGCTGTTTCCGATACTTAACAGTAGAGGAAAGCCTTGTAAACAAGGTGCTGTGTTGAG
TCTGTCTATTCAGTATATTCCAATGGAAAGAATGAGACTTTACCAAAAGGGTGTTGGTTTTGGTGTTGAATGTGTAGGAG
TTCCCGGTACATACTTCCCTTTAAGGAAAGGCGGTAGGGTTACTCTTTATCAGGATGCTCATGTTGATGACGGTACTCTT
CCGAGTGTACATCTTGATGGTGGGATTCAGTATAGACATGGAAAATGCTGGGAAGATATGGCTGATGCGATACGACGGGC
AAGGAGGTTGATTTATATCACAGGTTGGTCAGTTTTCCATCCGGTTAGGCTGGTTCGTCGTAACAATGATCCGACCCAAG
GTACATTAGGAGAGTTGCTTAAAGTCAAATCTCAAGAAGGTGTTAGAGTGTTGGTTTTGGTGTGGGATGATCCAACTTCA

**Figure S1. CRISPR target position and sequence in 12 *PLD*s.** Partial genomic DNA sequence of *PLD* is shown downstream from the start codon with introns underlined. CRISPR target sequence for sgRNA is highlighted in red.

PLDδ

ATGGCGGAGAAAGTATCGGAGGACGTTATGCTTCTACACGGTGACCTCGATTTGAAAATTGTTAAGGCTAGGAGGTTACC
TAACATGGATATGTTCTCAGAACATTTGCGCCGTCTCTTCACCGCCTGTAACGCCTGTGCTAGACCCACCGATACCGATG
ATGTCGATCCCAGAGATAAAGGCGAATTCGGTGATAAAAACATCCGTAGCCACCGTAAGGTTATCACCAGCGATCCTTAC
GTCACCGTCGTCGTTCCTCAAGCGACTCTAGCTCGAACACGTGTTTTGAAAAACTCACAAGAGCCTCTTTGGGACGAGAA
ATTCAACATTTCTATAGCGCATCCGTTTGCTTATCTCGAGTTCCAGGTCAAAGACGACGATGTTTTCGGTGCTCAAATCA
TTGGCACGGCTAAGATCCCGGTTCGAGACATCGCATCGGGAGAACGCATTTCTGGTTGGTTTCCTGTACTCGGTGCTTCG
GGAAAAACCGCCTAAGGCAGAAACTGCTATTTTCATCGATATGAAATTTACTCCGTTTGACCAGATCCATAGCTACCGATG
TGGAATCGCCGGAGATCCGGAGCGTAGGGGTGTTAGACGGACTTATTTCCCTGTGAGGAAAGGAAGTCAGGTGAGGCTTT

PLDε

ATGGAGCTTGAAGAACAGAAGAAGTACTTTCATGGAACACTAGAGATTACAATCTTCGATGCAACACCTTTTTCTCCTCC
ATTTCCTTTCAATGTAAGTCTATATCATATACTGAATTAGTGACTCTGATGATGTTATCATTGTGTTCATTTTCATGAGC
TTTTGTTATCTTTGTTCACAGTGTATATGTACAAAGCCAAAAGCAGCTTACGTGACCATCAAGATAAACAAGAAGAAAGT
TGCTAAAACAAGCTCAGAATACGACCGTATTTGGAATCAGACTTTTCAGATCCTCTGCGCTCACCCGGTTACTGACACGA
CTATCACCATCACGCTCAAGACTCGATGTTCTGTTCTCGGAAGATTCCGAATCTCTGCAGAACAGATCCTGACATCTAAT
TCAGCAGTTATCAACGGGTTCTTCCCTCTGATTGCAGACAATGGGTCAACGAAACGTAACTTGAAGCTTAAGTGTTTGAT
GTGGTTCAGACCGGCTTATCTTGAACCGGGATGGTGCAGAGCGCTTGAAGAAGCGTCTTTTCAAGGTATTAGGAACGCAA
GCTTCCCTCAAAGATCGAACTGCAGAGTCGTACTGTATCAAGATGCACACCACAAGGCCACATTTGATCCAAGGGTTGAT

PLDζ1

ATGGCATCTGAGCAGTTGATGTCTCCCGCCAGTGGTGGTGGACGCTACTTTCAGATGCAGCCTGAGCAATTTCCTTCGAT
GGTCTCTTCGCTCTTCTCTTTCGCGCCGGCTCCTACGCAGGAGACTAATCGTATTTTTGAAGAATTACCAAAAGCAGTGA
TCGTCTCTGTCTCTCGCCCTGATGCCGGCGATATTAGCCCTGTACTCTTGTCTTACACCATTGAGTGCCAATACAAGCAG
GCGAGTCCATCTCTTCTCTTTAGTAGGTCTTACTAATATCGTCGCATAATAGATGCTTGGATCAGAATCACTTAAGAGGA
GATGGGTATCAACTGTAAATTGTCATGTCTGACTATTGATCGAAGTAGTTCTTTTGGATACCTAGCTTGAATCTGATACC
ACTGATCGATGTTATATGTCCTTTGGAAGCCCGGAACAGAGACATATTTATCAATATTTATATGTTTCACCTTAGTTTAG
TGGAAAGGCTGCTTATTATAGTGGAGCTTTCGACTTTTGAATACTTGAGTCCTAACTTTAGCGCTTTTATGTAACAGTTC
AAGTGGCAGCTTGTGAAGAAAGCATCTCAAGTCTTTTATTTGCATTTTGCATTGAAGAAACGTGCTTTTATTGAAGAAAT
TCACGAGAAGCAGGAACAGGTTTGTGGTGACTTGATATATCAAGGAATTTTCTTCTACTAGTTTGTGATTCCCTTTGAGT
GAATATTTTAAAGGTTTTTCCTTGGTCTGTTGAATTGAGTTTGATGTGGTTTTCTTTCGAATATCACATTAAAGGTTAAA
GAATGGCTTCAAAATCTAGGAATAGGGGATCATCCACCCGTTGTGCAAGATGAAGATGCTGATGAAGTTCCGCTACATCA
AGATGAAAGTGCCAAAAATAGGTCACACTTGCTGGCATTTTTTTTTGCTTGGATAAGTTGCTGATGTGAGTCTATACTCT
TCCAACAGATGCCATCATTTTCTGACAAGTAGACTGTTTTGTTCAGAGATGTTCCTTCGAGCGCTGCTTTGCCAGTCATT
CGTCCTTTGGGAAGACAGCAGTCCATATCAGTTAGAGGAAAGCATGCAATGCAAGAATATCTGAATCATTTTCTGGGGAA
TCTTGATATCGTCAATTCACGGGAGGTATATTTGTAAAATCTATTTCACTTGGTTGTTCAACTACTTTAAAATTGTTAAT
ACCTAGTTATGTCAAGGGTTGTTTTCTCTTCTATTCTAGCTAGAAATCTTAGTACAGCAAGTGACTATGCGAAAATTTCC
ATGCTTTTCAAACTTGTATTCCTGTACACATTGATCGTCATTCCATTTCCGAGGCTATATGCATGCATAGTTAAATGCTA
AATGCCTAATATAGGTTTGCAGGTTTTTGGAGGTTTCGATGTTGTCATTCTCACCAGAGTATGGGCCCAAATTGAAAGAA
GACTATATCATGGTAAAACATCTACCGAAGTTTTCAAAGAGTGATGATGATTCTAATAGATGTTGTGGGTGCTGTTGGTT
CTGTTGCTGCAACGATAATTGGCAAAAGGTATGAAAATTCTCATTTTCTTGCTCATACTCGCGTACTATGTATTTTGCTA
TGAAAAATTGGTTTACCTCTGGAGTTAACTCTGGGATGACGTGTATGCTTTGTTTATGTGTTCTTTTGTTATTCTTGTCC
GCTCTCCTACTCTTGCTTTGTCTCTGAACTCGTTGAATGCCATGAATAGCTTATGTATCAGAAGAAACAAAATTCATCTT
TAGTAGTTCTTTTGTAATGATTTTTTTTTGTTACTACTCTCACTGGAGAGGTGTTCTGAAACAAACTTATCTGAATGCTT
CACCACCGTTGGGAGATATTAAGTTGATGAAGTAAATTGTTTTTCTAACTCTACTTTAGATCCATATGTTCTTCAATGTA
TGTGCCATCGAATACTGTTGTCTCCTTTTATAACTTCTGTCTTTGCTTCTGATCAGACATTCTAAATTAATGTGACGACA
AGATGGTATCTAAGGTTTCTTTTTCTCCAGTACAGGTGTGGGGGGTACTAAAGCCAGGTTTTCTCGCCTTATTGGAAGAT
CCATTTGATGCGAAGCTATTAGATATAATTGTTTTTGATGTCCTACCAGTTTCTAATGGAAATGATGGTGTGGATATATC
ACTAGCAGTAGAACTGAAGGATCATAATCCTTTGCGGCATGCATTCAAGGTCAGCAATTACTGTCCTACTTATCTTTTAA
TCAATTATATGTGTGCTTACTCCATCTAAATAAGCTTTCTTGATCATCTTTTTATACAATCCTCCTTGTTTGATAATCTA
CGATGTTGTTCTTTTATGCTTCATTGCTTCTTAGAGAAATTAAAATAAATTGTCCAAGTTATATATCCGAATTTTATTTC
TATTACGCATGGACACTAATGCGCCGTGGAGAACTGATAACCAAGAAAATTAGAATATTCCTTTTGTTGTAAATTAAAAT
GGGAGCCTACTGAATCCTTCTTTTATTAGCTAAAAGTTGAATCATCCTCATTTTTCTTGCCCCTCTTCTGCTGCACAAAT
ATTTCTGGGGTTATATGTCAAGAAAGCTATGGATAATCCATGTATATAACTTTTGTTAAAATATCCCAGGTAACATCTGG
AAACCGGAGTATAAGAATAAGGGCAAAGAATAGTGCAAAAGTTAAAGATTGGGTGGCTTCTATTAACGATGCTGCTCTTA
GACCTCCTGAGGGTTGGTGCCATCCCCATCGCTTTGGCTCATATGCTCCGCCGAGGGGTTTGACGGATGACGGAAGTCAA
GCCCAGTGGTTTGTAGATGGTGGAGCAGCTTTTGCAGCCATTGCTGCAGCGATTGAAAATGCTAAATCTGAGGTCTACCA
CTTTTCATTACTGGGTGTGGAGACTGATGATGTGTTGAATGTTATATATTTTCAACTGATCTGATTTGCCTTATAGATTT
TCATCTGTGGCTGGTGGGTGTGCCCAGAACTCTATCTTAGGCGTCCTTTTGACCCGCATACTTCATCCAGACTTGATAAC
TTGTTGGAGAATAAAGCTAAGCAAGGAGTTCAGGTACTTACTACAGAACATGCATGTACTTAATAGCTATTATCTTCTGT
TGTGAAGCATGTCTGTAGTCTGCAATGGAGATCTGACTTGTCAGGTTTGGTGGATTACTATCTTTCAACTTTTTCATTTT
TATTACTTTCTTTTTCAGATATACATCCTTATCTACAAGGAGGTTGCTCTTGCTTTAAAGATCAACAGTGTATATAGCAA
ACGCAGGCTTCTTGGCATTCATGAGAATGTGCGGGTACTTCGTTATCCTGATCATTTCTCAAGTGGTGTCTACCTCTGGT

PLDζ2

ATGTCGACGGATAAATTACTACTTCCTAACGGCGTTAAGTCAGACGGAGTCATCAGAATGACCAGAGCTGATGCTGCGGC
GGCGGCAGCTTCTTCTTCTCTCGGCGGTGGAAGTCAAATATTCGACGAGCTTCCCAAGGCTGCGATCGTCTCGGTCTCGA
GACCTGACACCACCGATTTTAGTCCCTTGCTTCTTTCTTACACCTTGGAGCTTCAGTATAAACAGGTATATATATAGTTT
ACAATGTCTGACTTGTAGTTTTAAAAGAACTTGATTTCCTCTCTGTTACATTGTGATTAGGGTTTTGATTTTGTCTATGT
ATACGAAATTTATGACAAAGAGATGCCATTAAGAGATTCAATTTCTTGCTAAAGAAGTTTATTCATGCATATATGAAATT
TCTAGTGGTTGTTTCTCTAAAAGTTTATGTTCTGTTTTTATTAGTAACAAACAAAGACCTTTTTTAATTGGTATTCTCTG
TTTTTGGTATATGTATAGTTCAAGTGGACATTACAAAAGAAGGCTTCTCAAGTTCTGTACTTACATTTTGCGTTGAAGAA
ACGTTTGATCATTGAAGAACTTCACGACAAGCAAGAACAGGTCAGAATGTTCAGGCTTTGCAGATTTGCTACTAACACAA
GCCAAACAAATAATTATTGAACTCCAGAGATGCGGTTTTCGACAGGTTAGAGAGTGGCTACACAGCTTGGGGATTTTTGA
TATGCAAGGATCAGTTGTGCAAGATGATGAAGAACCTGACGATGGTGCTCTTCCTCTGCACTATACTGAAGATAGTATCA
AGAACAGGTGCTTTTAAATGCCATAAAACTTTTCGGTTTTGTGGATTCAAAGTAAAAATTGTTTCGTTTAAATTTCTGAA
ACAGTAGCTATTTATGTTATGCAGGAATGTTCCTTCCCGTGCAGCGCTTCCAATCATTCGTCCAACGATAGGCCGGTCAG
AGACAGTTGTAGATCGTGGGAGAACCGCAATGCAAGGCTACTTGAGTCTCTTTCTAGGGAACTTGGACATTGTAAACTCC

**Figure S1.** Continued

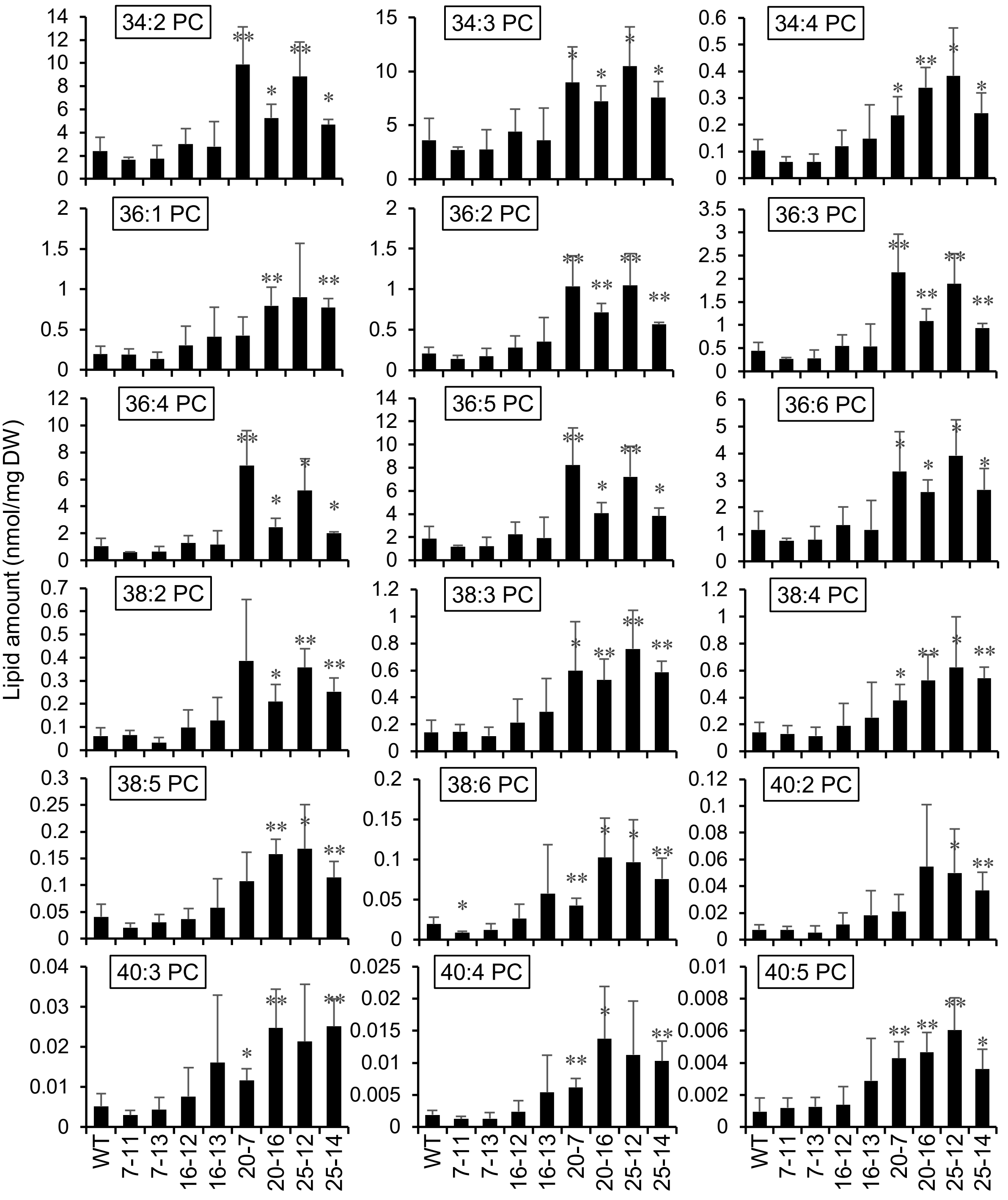


**Figure S2. PC levels in the selected T3 mutants.** Total lipids were extracted from the selected T3 mutants, and PC was separated and quantified by ESI-MS/MS. Data are shown here as the amounts of individual molecular species. Values represent mean ± S.D. Asterisks denote significant difference from WT as determined by Student's t-test (*$p<0.05$; **$p<0.01$; n=4 biological replicates). DW, dry weight of plant tissue.

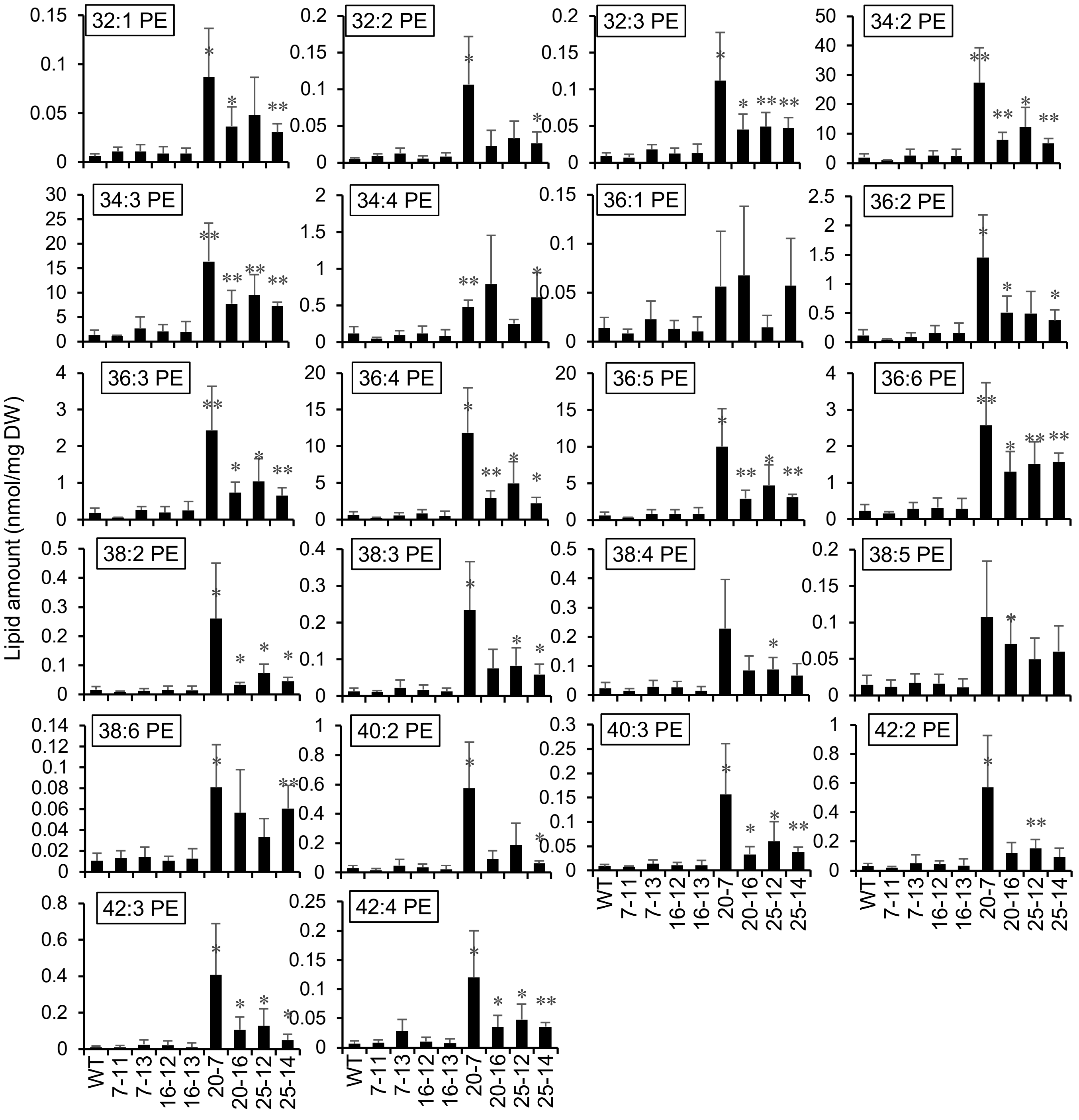


**Figure S3. PE levels in the selected T3 mutants.** Total lipids were extracted from the selected T3 mutants, and PE was separated and quantified by ESI-MS/MS. Data are shown here as the amounts of individual molecular species. Values represent mean ± S.D. Asterisks denote significant difference from WT as determined by Student's t-test (*p<0.05; **p<0.01; n=4 biological replicates). DW, dry weight of plant tissue.

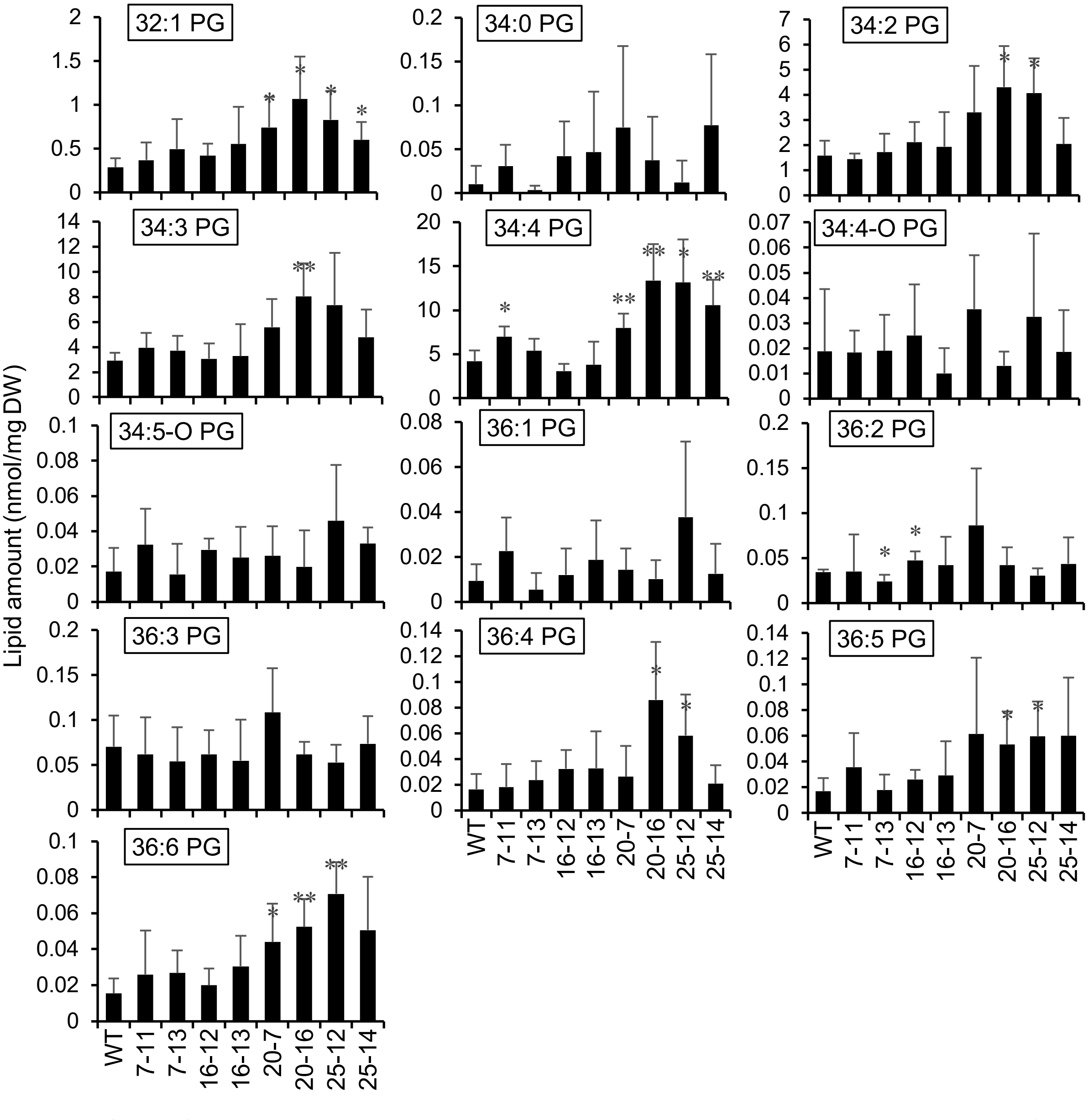


**Figure S4. PG levels in the selected T3 mutants.** Total lipids were extracted from the selected T3 mutants, and PG was separated and quantified by ESI-MS/MS. Data are shown here as the amounts of individual molecular species. Values represent mean ± S.D. Asterisks denote significant difference from WT as determined by Student's t-test (*p<0.05; **p<0.01; n=4 biological replicates). DW, dry weight of plant tissue.

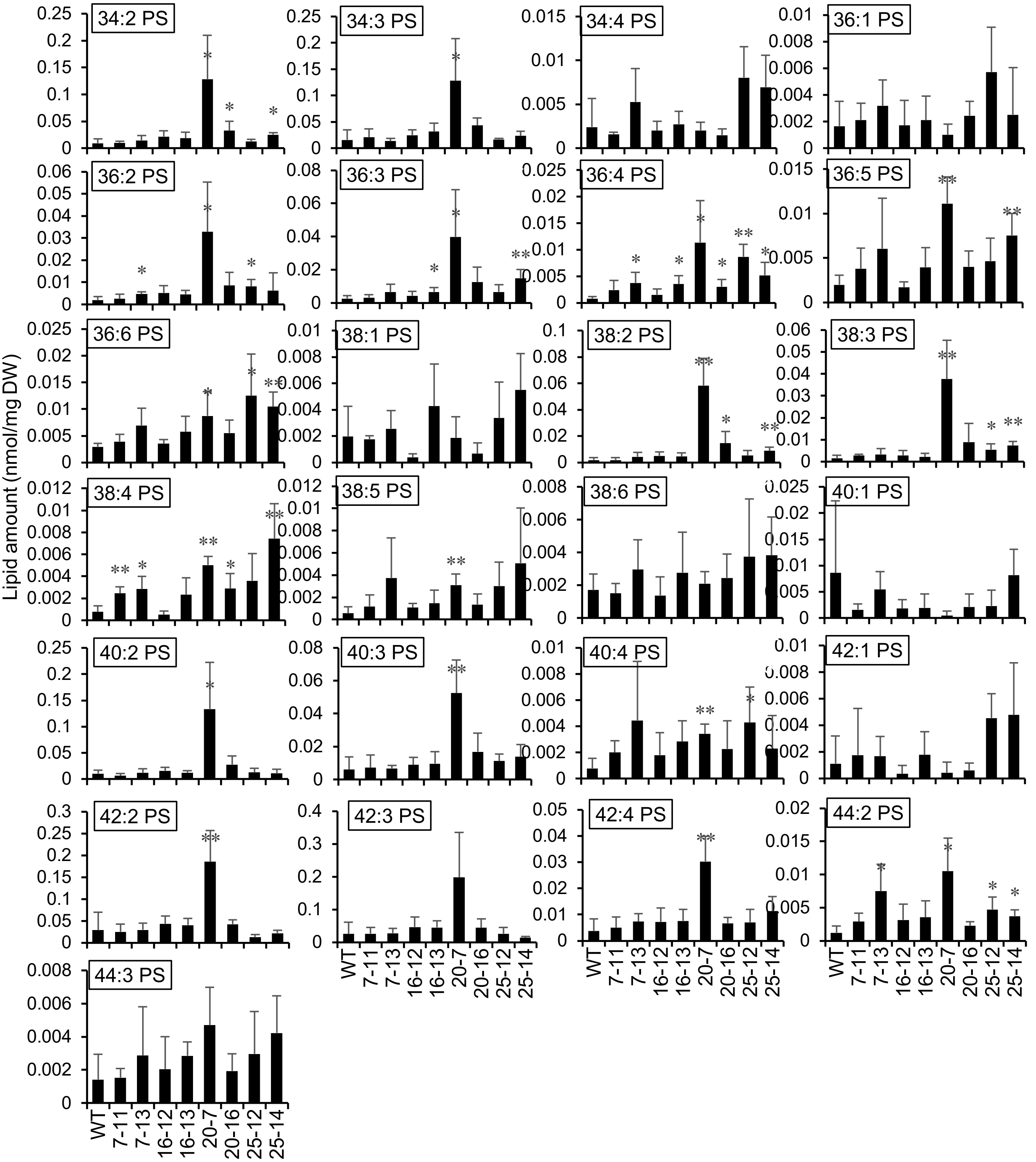


**Figure S5. PS levels in the selected T3 mutants.** Total lipids were extracted from the selected T3 mutants, and PS was separated and quantified by ESI-MS/MS. Data are shown here as the amounts of individual molecular species. Values represent mean ± S.D. Asterisks denote significant difference from WT as determined by Student's t-test (*p<0.05; **p<0.01; n=4 biological replicates). DW, dry weight of plant tissue.

### Table S1. Sequence of oligonucleotides used in this study (5'→3')

| Gene | Target site | Forward | Reverse |
|---|---|---|---|
| For sgRNA oligonucleotides | | | |
| Gene | Target site | Forward | Reverse |
| *PLDα1* | 1 | ATTGCCTAACACCACCACCATGG | AAACCCATGGTGGTGGTGTTAGGC |
| | 2 | ATTGCGGCGAGGAAGTGGATCAGT | AAACACTGATCCACTTCCTCGCCG |
| *PLDα2* | 1 | ATTGATCATCTCCATGCTGAAGG | AAACCCTTCAGCATGGAGATGATC |
| | 2 | ATTGAAGGATAAAAACTGGAATCG | AAACCGATTCCAGTTTTTATCCTT |
| *PLDα3* | 1 | ATTGTATAACTTGTCCTCCCGTGA | AAACTCACGGGAGGACAAGTTATA |
| | 2 | ATTGTTGAATGTGCGGTGTAGACG | AAACCGTCTACACCGCACATTCAA |
| *PLDβ1* | 1 | ATTGTAAGCACCAAAGTTACCTTG | AAACCAAGGTAACTTTGGTGCTTA |
| | 2 | ATTGCCTAGAAGTTCATCCACAGG | AAACCCTGTGGATGAACTTCTAGG |
| *PLDβ2* | 1 | ATTGCTGAGTAAATTTGTTCAACA | AAACTGTTGAACAAATTTACTCAG |
| | 2 | ATTGCGACTGATATGGACACGTA | AAACTACGTGTCCATATCAGTCGC |
| *PLDγ1* | 1 | ATTGGGAGCTGTTGGAATCCCAA | AAACTTGGGATTCCAACAGCTCCC |
| | 2 | ATTGTTTCCATAACAGACTTGGT | AAACACCAAGTCTGTTATGGAAAC |
| *PLDγ2* | 1 | ATTGTTCAGCAGCACTATGAGCAA | AAACTTGCTCATAGTGCTGCTGAA |
| | 2 | ATTGGGATTTCGTTTTCTGAGTT | AAACAACTCAGAAAACGAAATCCC |
| *PLDγ3* | 1 | ATTGCAAAAAACATCCCACCAACA | AAACTGTTGGTGGGATGTTTTTTG |
| | 2 | ATTGCAACTCTCCTAATGTACCT | AAACAGGTACATTAGGAGAGTTGC |
| *PLDδ* | 1 | ATTGATCTCCGGCGATTCCACAT | AAACATGTGGAATCGCCGGAGATC |
| | 2 | ATTGTCTAGCACAGGCGTTACAGG | AAACCCTGTAACGCCTGTGCTAGA |
| *PLDε* | 1 | ATTGTTGGATCAAATGTGGCCTTG | AAACCAAGGCCACATTTGATCCAA |
| | 2 | ATTGACATCGAGTCTTGAGCGTGA | AAACTCACGCTCAAGACTCGATGT |
| *PLDζ1* | 1 | ATTGAGCATCTTCATCTTGCACAA | AAACTTGTGCAAGATGAAGATGCT |
| | 2 | ATTGTTGGCATTCATGAGAATGTG | AAACCACATTCTCATGAATGCCAA |
| *PLDζ2* | 1 | ATTGACTCAAGTAGCCTTGCATTG | AAACCAATGCAAGGCTACTTGAGT |
| | 2 | ATTGAATGACCAGAGCTGATGCTG | AAACCAGCATCAGCTCTGGTCATT |
| For PCR of genomic DNA | | | |
| Gene | | Forward | Reverse |
| *PLDα1* | | TGTGTTCTGGTCAATTCTTATT | GAGAAGAATGTGTATGGCACT |
| *PLDα2* | | GTGAATTTACGTATTTTGGTAATC | GACAAAGTTTCCAGGGATATG |
| *PLDα3* | | CAAAAGACTAACGGACTCAT | CTCTAACTAATGTCACGTCG |
| *PLDβ1* | | GACCATTAGACTATAGCCAC | ATGAGAGTTACTTGGAAACG |
| *PLDβ2* | | ATATTTGGGTTTCTTGTGCTAA | ATTGAAGTATACTGAATCGACAA |
| *PLDγ1* | | CGGCTTATACTGAGACTATG | CAACACCCATTTGGTAAAGT |
| *PLDγ2* | | GATGGTTGAATTGTTACATGG | AATAGACAGACTCAACATAGC |
| *PLDγ3* | | GATACAAGTAGTGGCTCTTT | AAGACTCCTTGAAGTTGGAT |
| *PLDδ* | | AAAGTATCGGAGGACGTTAT | TTCCCGTTATCTAACCCAAT |
| *PLDε* | | TACGTGACCATCAAGATAAAC | GTACAAGAACAAGATTAGGGT |
| *PLDζ1* | | AGAAAGCATCTCAAGTCTTTT | ATGACGAGTTTTTCATGGTG |
| *PLDζ2* | | GAGTATTTCGGACATGCAAT | GTGACATACCCTTCTTTCAT |
| For DNA sequencing | | | |
| Gene | | Target site 1 | Target site 2 |
| *PLDα1* | | GTTGTATGCGACGATTGATCTGC | GCAGATCAATCGTCGCATACAAC |
| *PLDα2* | | CCCAAAGTGGTTTGAGTCTTTTC | GAAAAGACTCAAACCACTTTGGG |
| *PLDα3* | | CCAAACTCCATGTACGTGTGAAG | CTTCACACGTACATGGAGTTTGG |
| *PLDβ1* | | GCCTCCTCACTATTCGTATCAAG | TACATTATCATGACCTGTTCCCC |
| *PLDβ2* | | GGTGCTGTCATAGGAAGAACATATG | CAATTCCTATAAGCTGTGAACCG |
| *PLDγ1* | | GGTTAGGTCGGAAGAAAGTGG | TTCAGCAGCACTATGAGCAAC |
| *PLDγ2* | | CGGAGGATTCGTAAAGTGGATG | TACATCGAAATGCTGCATCCAC |
| *PLDγ3* | | CTTGTAAACAAGGTGCTGTGTTG | CAACACAGCACCTTGTTTACAAG |
| *PLDδ* | | CTATAGCGCATCCGTTTGCTTATC | GATAAGCAAACGGATGCGCTATAG |
| *PLDε* | | GCTTAAGTGTTTGATGTGGTTCAG | CTGAACCACATCAAACACTTAAGC |
| *PLDζ1* | | CTTCATCCAGACTTGATAAC | GAATTGACGATATCAAGATTC |
| *PLDζ2* | | GGCTTCTCAAGTTCTGTACTTAC | GTAAGTACAGAACTTGAGAAGCC |
